# *stk32a* links sleep homeostasis to suppression of sensory and motor systems

**DOI:** 10.1101/2025.09.09.675098

**Authors:** Steven Tran, Jasmine Emtage, Chaodong Zhang, Xiaoyao Liu, Marina Lecoeuche, Andrey Andreev, Grigorios Oikonomou, Sujatha Narayan, Brianna Garcia, Tasha Cammidge, Cristina Gonzales, Hannah Hurley, Misha Yap, Shan Li, Feng Wang, Ting-Yu Wang, Misha B. Ahrens, Tsui-Fen Chou, Min Xu, Qinghua Liu, David A. Prober

**Affiliations:** Tianqiao and Chrissy Chen Institute for Neuroscience, Division of Biology and Biological Engineering, California Institute of Technology, Pasadena, CA, 91125, USA; New Cornerstone Science Laboratory, National Institute of Biological Science, Beijing (NIBS), Beijing 102206, China; Tsinghua Institute of Multidisciplinary Biomedical Research (TIMBR), Tsinghua University, Beijing 10206, China; Institute of Neuroscience, Center for Excellence in Brain Science and Intelligence Technology, Chinese Academy of Sciences, Shanghai 200031, China; Janelia Research Campus, Howard Hughes Medical Institute, Ashburn, VA 20147, USA

**Keywords:** Sleep homeostasis, Kinase, Serotonin, Raphe, Neurotensin, Zebrafish

## Abstract

Sleep is regulated by a homeostatic process and associated with an increased arousal threshold, but the genetic and neuronal mechanisms that implement these essential features of sleep remain poorly understood. To address these fundamental questions, we performed a zebrafish genetic screen informed by human genome-wide association studies. We found that mutation of *serine/threonine kinase 32a* (*stk32a*) results in increased sleep and impaired sleep homeostasis in both zebrafish and mice, and that *stk32a* acts downstream of neurotensin signaling and the serotonergic raphe in zebrafish. *stk32a* mutation reduces phosphorylation of neurofilament proteins, which are co-expressed with *stk32a* in neurons that regulate motor activity and in lateral line hair cells that detect environmental stimuli, and ablating these cells phenocopies *stk32a* mutation. Neurotensin signaling inhibits specific sensory and motor populations, and blocks stimulus-evoked responses of neurons that relay sensory information from hair cells to the brain. Our work thus shows that *stk32a* is an evolutionarily conserved sleep regulator that links neuropeptidergic and neuromodulatory systems to homeostatic sleep drive and changes in arousal threshold, which are implemented through suppression of specific sensory and motor systems.

## INTRODUCTION

Sleep is regulated by a homeostatic process, in which sleep need increases with time spent awake, and is associated with an increased arousal threshold, such that a stronger stimulus is required to arouse a sleeping animal compared to an animal that is awake ^1^. While several molecular and cellular processes have been proposed to underlie sleep homeostasis ^2^, the genetic and neural circuit mechanisms through which homeostatic sleep pressure is implemented in the brain remains poorly understood, particularly in vertebrates. It is also unclear how changes in arousal threshold are linked to sleep homeostasis, sensory input, and motor output. Genetic screens in invertebrates ^3^ and vertebrates ^4^ have been a useful approach to identify genes and mechanisms that underlie sleep. However, the relevance of genes identified in these model organisms to human sleep can be unclear. Genes that regulate human sleep can be identified by studies of families with highly-penetrant sleep phenotypes ^5^, but this approach is limited by the rarity of these phenotypes. Alternatively, one can identify genes relevant to human sleep based on genome-wide association studies (GWAS) that identify common genetic variants associated with specific sleep phenotypes. Indeed, recent GWAS have identified >100 genomic loci containing variants associated with several sleep traits ^6^. However, these variants are non-coding, and most loci contain several nearby genes. As a result, for most loci the causal gene is unknown, and candidate genes must be experimentally validated using an animal model.

Here we describe a genetic screen that identified *stk32a* as a key regulator of both homeostatic sleep drive and arousal threshold. Using genetics, pharmacology, optogenetics, cell ablation, phosphoproteomics, and calcium imaging, we identify genetic and neuronal mechanisms through which *stk32a* regulates sleep. More broadly, we show that homeostatic sleep pressure is implemented by suppression of specific sensory and motor systems, and uncover a mechanism that links homeostatic sleep pressure with increased arousal threshold.

## RESULTS

### A zebrafish genetic screen identifies *stk32a* as a novel sleep regulator

We selected 30 genes in 23 genomic loci that were associated with self-reported sleep traits among UK Biobank subjects ^7,8^ for validation in zebrafish. These genes were selected because they had not been implicated in sleep, and there was no evidence they are required for development. Due to the presence of multiple zebrafish paralogs for some human genes, we identified 43 zebrafish orthologs of the 30 human genes. We used CRISPR/Cas9 to induce an insertion/deletion (indel) mutation in each zebrafish gene, resulting in a predicted null allele due to a shift in the translational reading frame and premature stop codon. Using a high-throughput sleep assay ^9^, we assessed the impact of each mutation on zebrafish behavior at 5 days post-fertilization (dpf), identifying a statistically significant sleep and/or locomotor activity phenotype for 20 of the 43 mutants **(Extended Data Fig. 1a)**.

Several single nucleotide polymorphisms (SNPs) in intronic regions of the human *STK32A* gene are associated with daytime napping **(Fig. 1a)** ^8^. Since the SNPs might be linked to the neighboring genes *DPYSL3* (3’ to *STK32A*, **Fig. 1a)** or *PPP2R2B* (5’ to *STK32A*, not shown), we mutated the single zebrafish orthologs of *STK32A* **(Extended Data Fig. 1b)** and *DPYSL3*, and the two zebrafish paralogs of *PPP2R2B* (*ppp2r2ba* and *ppp2r2bb*). There was no significant change in sleep or locomotor activity for *dpysl3*, *ppp2r2ba*, or *ppp2r2bb* mutants compared to their sibling controls **(Extended Data Fig. 1c-e)**.

**Figure 1.**
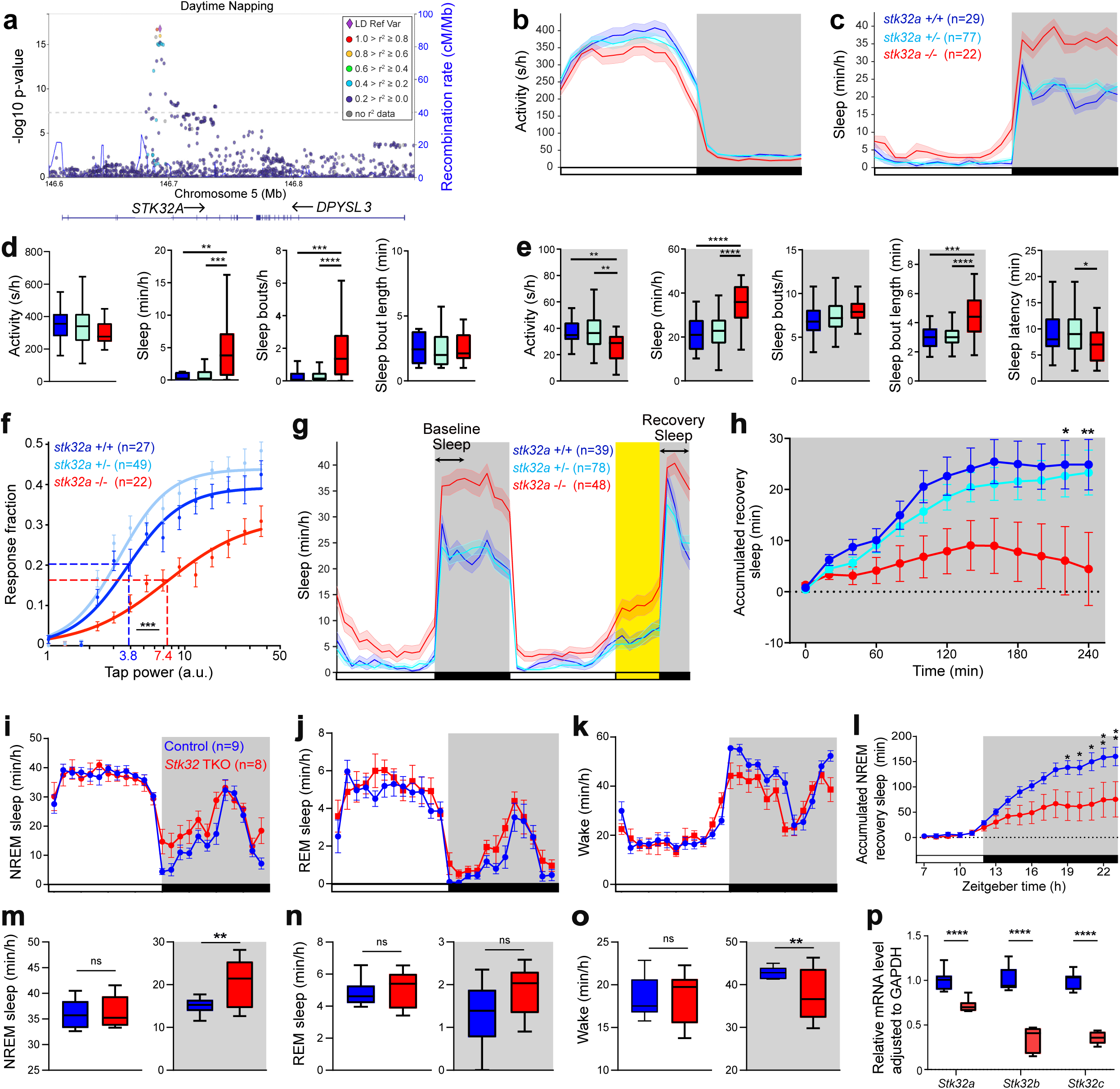
*stk32a* −/− zebrafish and *Stk32* TKO mice show increased sleep and defective sleep homeostasis. **(a)** SNPs within the human *STK32A* locus that show genome-wide association with variation in daytime napping. The lead SNP (rs10875622) is shown in purple and additional SNPs are colored according to their correlation with the lead SNP, based on CEU HapMap haplotypes. Dashed line represents genome-wide significance threshold (P<5×10^-8^). **(b-h)** Zebrafish experiments. Locomotor activity **(b)** and sleep **(c)** during the 5th day and night of development. Lines and shading represent mean ± SEM. **(d,e)** Box plots show locomotor activity, sleep, number of sleep bouts, and sleep bout length for day **(d)** and night **(e)**, and sleep latency at night. **(f)** Acoustic stimulus-response curves indicate an EP50 of 7.4 for *stk32a −/−* fish and 3.8 for WT siblings. Mean ± SEM responses for 30 trials at each stimulus intensity are shown. **(g)** Mean ± SEM sleep during a 48-hour period (5-6 dpf). Baseline sleep was measured during the first 4 hours of the first night (arrow in first gray box). Fish were sleep deprived for the first 6 hours of the second night by maintaining daytime white light illumination (yellow). White lights were then turned off and recovery sleep was measured for the last 4 hours of the night (arrow in second gray box). **(h)** Accumulated recovery sleep following sleep deprivation was reduced for *sk32a −/−* fish compared to controls. **(i-p)** Mouse experiments. *Stk32* triple adult brain chimeric knockout (TKO) resulted in increased NREM sleep **(i, m)**, no change in REM sleep **(j, n)**, and decreased wake **(k,o)** during the dark phase. **(l)** Accumulated NREM recovery sleep following sleep deprivation was reduced for TKO mice compared to controls. **(p)** *Stk32a, Stk32b* and *Stk32c* mRNA levels in TKO mice compared to controls. Black bars and dark shading indicate night. n = number of animals. ns, P>0.05; *P<0.05; **P<0.01; ***P<0.001; ****P<0.0001 by Kruskal-Wallis test with Dunn’s test **(d,e,h**), extra sum-of-squares F test **(f)**, and 2-way ANOVA with Sidak’s test **(l-p)**.

In contrast, *stk32a −/−* fish slept almost twice as much at night, with a small increase in sleep during the day, compared to sibling controls **(Fig. 1c-e)**. The increase in daytime sleep resulted from more sleep bouts **(Fig. 1d)**, indicating wake fragmentation. In contrast, the increased sleep at night was due to longer sleep bouts, indicating sleep consolidation, as well as a shorter sleep latency after lights out **(Fig. 1e)**. Locomotor activity during the day was unaffected in *stk32a* mutants **(Fig. 1b,d)**, suggesting the increase in sleep is unlikely due to sickness, developmental defects, or locomotor impairment. To distinguish sleep from quiet wakefulness, we tested the arousal threshold of *stk32a* mutants by delivering an acoustic stimulus over a range of intensities at night ^10^. We determined the fraction of fish that responded to each stimulus and calculated the stimulus power that resulted in the half maximal response (effective power 50 [EP50]). We observed a large increase in EP50 for *stk32a −/−* fish compared to sibling controls **(Fig. 1f)**, indicating that *stk32a* mutants have an increased arousal threshold.

### *stk32a* is required for homeostatic but not circadian regulation of sleep in zebrafish

Sleep is regulated by a homeostatic process, in which sleep pressure accumulates with time spent awake and dissipates with sleep, and a circadian process, which determines when most sleep occurs during the 24-hour circadian cycle ^1^. We explored the relationship between *stk32a* and circadian regulation of sleep by performing two experiments. First, we entrained *stk32a* mutants and their sibling controls to 14:10 hour light:dark cycles for 5 days, and then shifted them to either constant light or constant dark. Behavioral circadian rhythms were maintained in *stk32a −/−* fish in both cases **(Extended Data Fig. 2a-f)**, indicating that Stk32a is dispensable for circadian regulation of sleep. In addition, light was arousing and dark was sedating in *stk32a −/−* fish **(Extended Data Fig. 2a,b,d,e)**, indicating that Stk32a is not required for the masking effects of light and dark on behavior. Second, we raised and tested *stk32a* mutants in constant light, which prevents the establishment of entrained circadian rhythms ^11^. These fish lacked circadian patterns of locomotor activity and sleep, as expected, but *stk32a −/−* fish still slept more than controls **(Extended Data Fig. 2g,h)**. Thus, the *stk32a* mutant sleep phenotype is independent of entrained circadian rhythms.

We next asked whether *stk32a* is required for homeostatic regulation of sleep. To test this hypothesis, we maintained daytime white light illumination for the first 6 hours of the night, which results in a large decrease in sleep compared to the prior night **(Fig. 1g)**. We then turned the lights off for the remaining 4 hours of the night, and compared the amount of recovery sleep to the prior night of baseline sleep ^12^. Control fish showed a large increase in recovery sleep for the first few hours after sleep deprivation compared to the prior night, whereas *stk32a* mutants showed no significant increase **(Fig. 1h)**. This result indicates that *stk32a* mutants have impaired rebound sleep following sleep deprivation, thus implicating *stk32a* in homeostatic regulation of sleep. This observation, together with the elevated baseline sleep in *stk32a* mutants, is consistent with the hypothesis that *stk32a* mutants have elevated, and nearly saturated, homeostatic sleep pressure, similar to mice that express a constitutively active form of CaMKII ^13^.

### *Stk32a/b/c* mutant mice phenocopy zebrafish *stk32a* mutants

Similar to humans, mice have 3 paralogs (*Stk32a*, *Stk32b*, and *Stk32c)* of the zebrafish *stk32a* gene. To determine whether these mouse genes regulate sleep, we used an adult somatic CRISPR/Cas9 mutagenesis approach (adult brain chimeric knockout (ABC-KO) ^14^), which avoids compensatory mechanisms that may arise due to loss of gene function during development. The mRNA level of each gene was significantly, though not fully, reduced **(Fig. 1p and Extended Data Fig. 3d)**. *Stk32a* ABC-KO mice did not exhibit sleep defects compared to controls **(Extended Data Fig. 3a-g)**, possibly due to redundant functions of *Stk32b* and *Stk32c*. Indeed, *Stk32* triple ABC-KO (TKO) mice exhibited a significant increase in NREM sleep during the dark phase due to a trend towards more sleep bouts **(Fig. 1i,m, and Extended Data Fig. 3i)**, without changes in REM sleep **(Fig. 1j,n and Extended Data Fig. 3k,l),** and a reduction in time spent awake due to shorter wake bouts **(Fig. 1k,o and Extended Data Fig. 3n),** indicative of wake fragmentation. To test if the mouse *Stk32* paralogs play a role in sleep homeostasis, we sleep deprived mice and quantified NREM recovery sleep. Although *Stk32a* ABC-KO mice had normal rebound sleep **(Extended Data Fig. 3h)**, *Stk32* TKO mice had significantly reduced rebound sleep compared to controls **(Fig. 1l)**, demonstrating that the *Stk32* paralogs regulate sleep homeostasis in mice, similar to zebrafish **(Fig. 1h)**. These experiments combined with the GWAS data suggest that *stk32a* regulates sleep in zebrafish, mice, and humans.

### Pharmacological inhibition of Stk32a promotes sleep in zebrafish and mice

As an alternative approach to test the role of Stk32a in sleep, we acutely inhibited Stk32a by treating wild-type (WT) zebrafish with the small molecules TAE684 and SB-245391, which inhibit human Stk32a, Stk32b, and Stk32c kinase activity ^15,16^. Both drugs increased sleep and decreased locomotor activity during the day and night, due to more sleep bouts during the day and longer sleep bouts at night **(Extended Data Fig. 4a,c)**, similar to *stk32a −/−* fish. Treatment with either inhibitor did not enhance the *stk32a −/−* sleep phenotype **(Extended Data Fig. 4b,d)**, suggesting that the drug-induced sleep phenotypes are due to specific inhibition of Stk32a. Alternatively, there may be a physiological ceiling effect, with *stk32a −/−* fish sleeping so much at night that the phenotype cannot be enhanced by other sleep-inducing perturbations. However, this is unlikely to be the case because overexpression of the sedating neuropeptide VF (NPVF)^17^ in *stk32a* −/− fish increased sleep compared to both *stk32a* −/− fish and *stk32a* +/− fish that overexpressed NPVF **(Extended Data Fig. 9b)**. We next asked if acute inhibition of the mammalian Stk32 proteins also promotes sleep by treating WT mice with TAE684 during the dark phase, when the *Stk32* TKO phenotype was observed. Drug treatment resulted in a large increase in both NREM and REM sleep due to more sleep bouts **(Extended Data Fig. 3o,p)**, and less time spent awake due to more but shorter wake bouts **(Extended Data Fig. 3q)**, indicative of wake fragmentation. These results show that both chronic genetic loss and acute pharmacological inhibition of Stk32a in zebrafish and Stk32a/b/c in mice results in increased sleep.

### *stk32a* is expressed in sensory, neuropeptidergic, and motor systems in zebrafish and mice

To explore how *stk32a* might regulate sleep, we next characterized its expression pattern by performing hybridization chain reaction (HCR) on 6-dpf zebrafish, which revealed that *stk32a* is expressed in three types of cells. First, *stk32a* is expressed in sensory cells, including hair cells in lateral line neuromasts **(Fig. 2d,e)**, which are sensory organs that detect external stimuli such as water flow and vibrations, and the inner ear (**Fig. 2d,e)**. *stk32a* is also expressed in the inner nuclear layer of the retina **(Fig. 3f)**, and in the olfactory epithelium **(Fig. 2a)**. Second, *stk32a* is expressed in GABAergic neurons in the neurosecretory preoptic area of the hypothalamus (NPO) **(Fig. 2b,i)**, subsets of which express the neuropeptides *neurotensin* (*nts*) and somatostatin (*sst1.1*) **(Fig. 2b,g,h)**. Third, *stk32a* is expressed in neurons that regulate motor functions, including the oculomotor nucleus (nIII) that controls eye movements **(Fig. 2b,j and Extended Data Fig. 5a)**, Mauthner neurons that regulate escape responses **(Fig. 2b,j)** ^18^, and cholinergic neurons in the spinal cord **(Fig. 2c,d and Extended Data Fig. 5c)** that express the fast motor neuron markers *neurofilament medium chain a* (*nefma*)^19^ **(Fig. 2k)**, *neurofilament medium chain b (nefmb), neurofilament light chain a (nefla),* and *neurofilament light chain b* (*neflb*)^19^ **(Extended Data Fig. 5e,g,i)**. *stk32a* is also expressed in a subset of cholinergic T-reticular interneurons in the hindbrain known as cranial relay neurons (CRNs) ^20^ **(Fig. 2c,j and Extended Data Fig. 5b)** that promote motor activity via interactions with the Mauthner neurons ^21,22^ and the nucleus of the medial longitudinal fasciculus (nMLF) ^23^. *stk32a* is also expressed in eurydendroid neurons in the cerebellum **(Fig. 2c,l)**, which regulate motor function ^24^. Together, these observations suggest that *stk32a* may play roles in sensory, neuropeptidergic, and motor systems in zebrafish.

**Figure 2.**
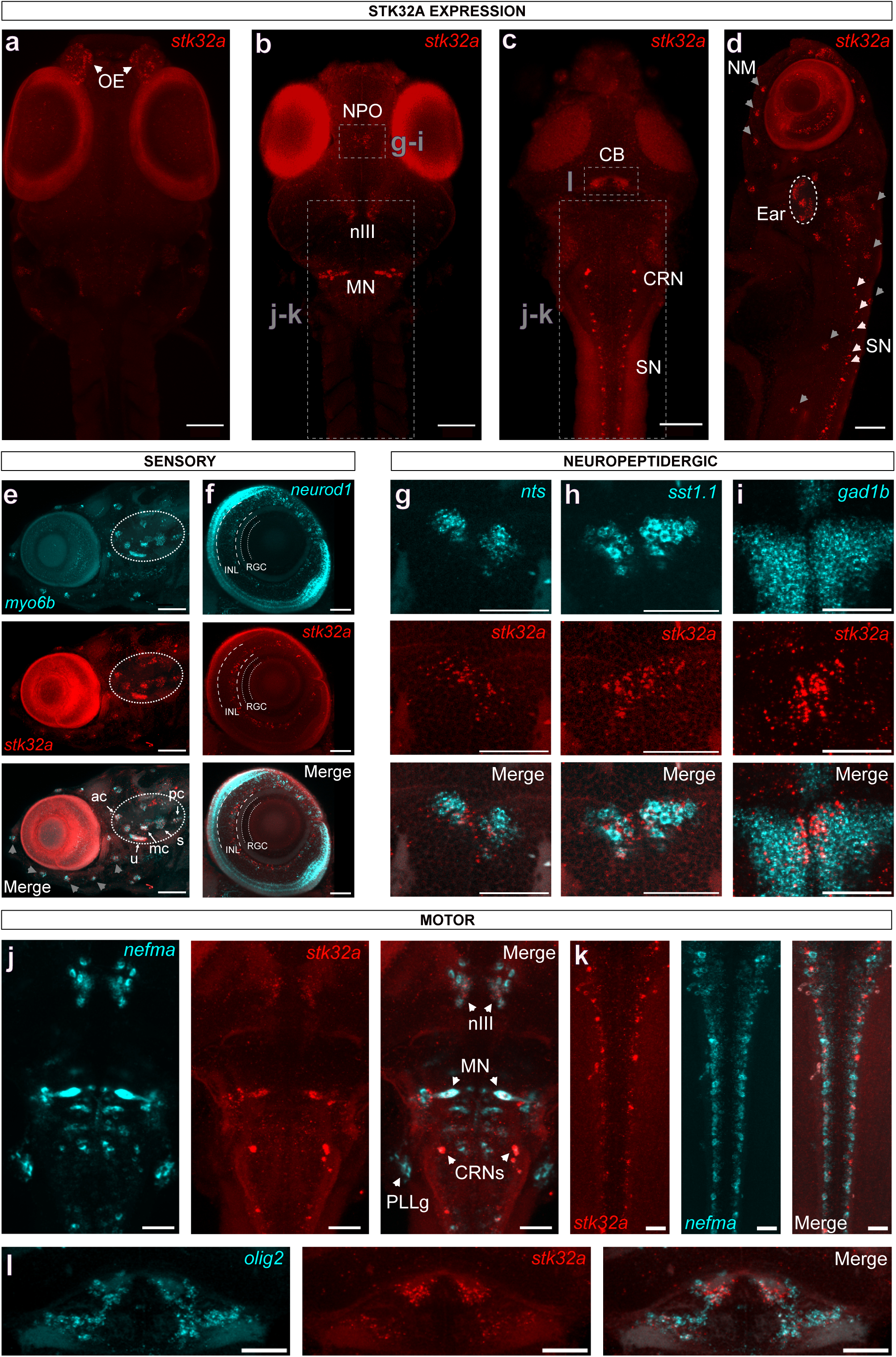
*stk32a* is expressed in sensory, neuropeptidergic, and motor regions in larval zebrafish. HCR in 6-dpf zebrafish reveals s*tk32a* expression in sensory regions including the olfactory epithelium (OE) (**a**), the inner ear (circled regions in **d,e**, including the five sensory patches: saccule (s), utricle (u), anterior cristae (ac), medial cristae (mc), and posterior cristae (pc)), neuromasts (NM) (gray arrows in **d,e**), and the inner nuclear layer (INL) of the retina **(f)**. Boxed regions in **(b,c)** are magnified in **(g-l**). **(g-i)** *stk32a* is expressed in GABAergic neurons in the neurosecretory preoptic area (NPO) **(i)** that co-express the neuropeptides *nts* **(g)** and *sst1.1* **(h)**. *stk32a* expression in motor systems includes the oculomotor nucleus (nIII) **(j)**, Mauthner neurons (MN) **(j)**, cranial relay neurons (CRNs) **(j)**, spinal cord neurons (SN) (**c,k,** and white arrows in **d**), and eurydendroid cells in the cerebellum (CB) **(l)**. Dorsal **(a-c, g-l)** and side **(d-f)** views are shown. Scale bars: 100 μm **(a-d)**, 50 μm **(e-j, l)**, or 25 μm **(k)**

**Figure 3.**
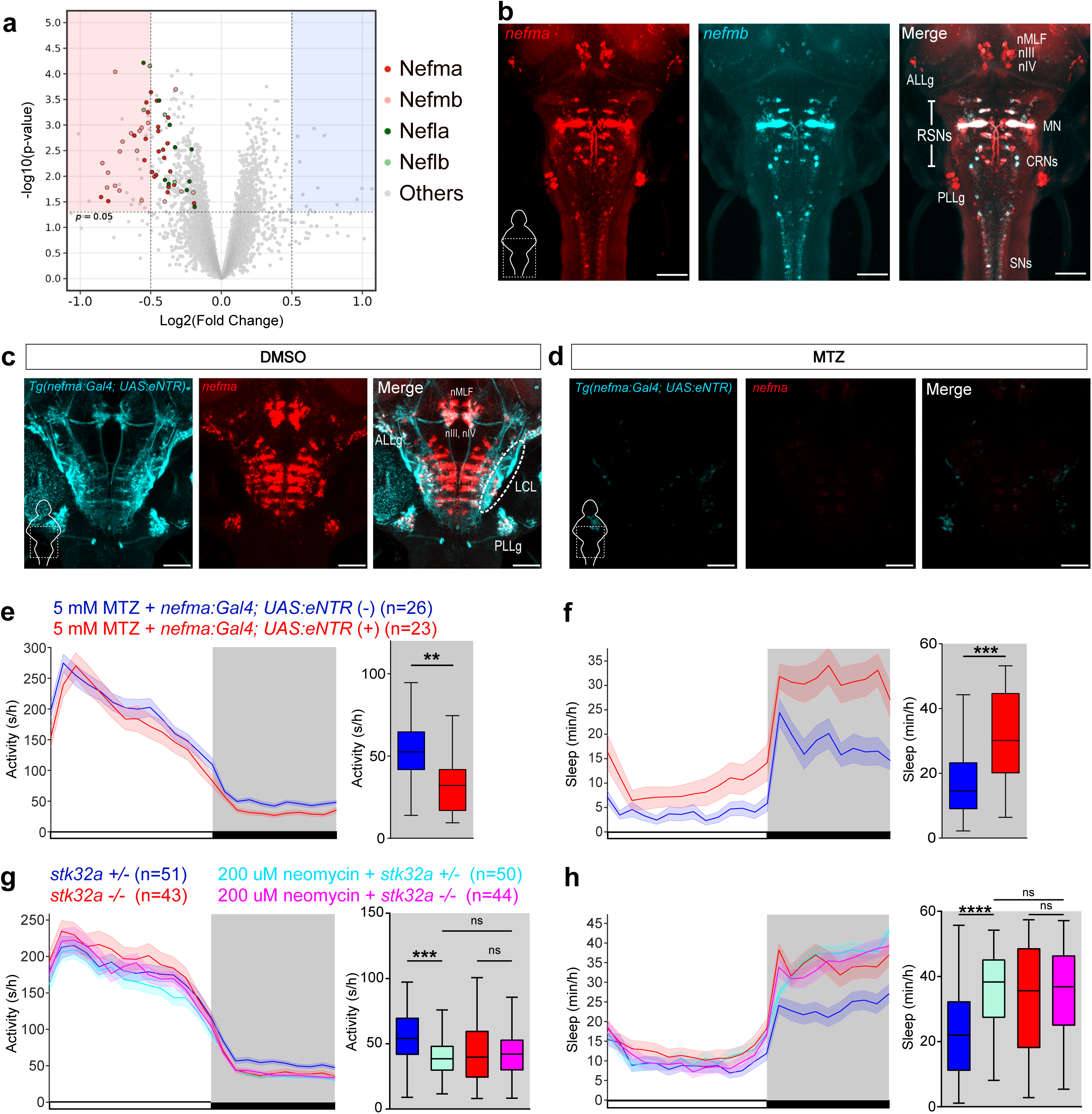
*nefma*-expressing neurons and lateral line neuromasts are required for normal sleep levels. **(a)** Volcano plot showing phosphopeptides that are less (red box) or more (blue box) abundant in the brains of *stk32a −/−* compared to *stk32a +/+* zebrafish, for changes that are least 1 fold and statistically significant (P<0.05, dashed lines). Nefma (red), Nefmb (orange), Nefla (dark green), Neflb (light green), and other phosphopeptides (gray) are shown. **(b)** HCR in 6-dpf zebrafish shows expression of *nefma* and *nefmb* in the anterior and posterior lateral line ganglia (ALLg and PLLg), cranial nerve nuclei (nIII, nIV), Mauthner neurons (MN), nucleus of the medial longitudinal fasciculus (nMLF), other reticulospinal neurons (RSNs), cranial relay neurons (CRNs) and spinal motor neurons (SNs). Note that *nefmb* expression is lower than *nefma* in the ALLg and PLLg, see Extended Data Fig. 5d for clearer *nefmb* expression. **(c,d)** *Tg(nefma:Gal4); Tg(UAS:eNTR-mYFP)* zebrafish treated with DMSO vehicle **(c)** or MTZ **(d)** showing *nefma* HCR and YFP immunostaining. The membrane-bound YFP (cyan) shows that *nefma* is expressed in lateral line ganglion neurons that innervate neuromasts and whose afferent projections form the lateralis central lintel (LCL, white dashed oval). Scale bars: 100 μm. **(e,f)** Locomotor activity and sleep of 5-dpf *Tg(nefma:Gal4); Tg(UAS:eNTR-mYFP)* (red) and non-transgenic sibling control (blue) zebrafish treated with MTZ. Genotypes indicate fish that were visually positive or negative for eNTR-mYFP fluorescence. **(g,h)** *stk32a +/−* fish treated with 200 μM neomycin (cyan) and *stk32a −/−* fish treated with vehicle control (red) show less locomotor activity and more sleep at night compared to vehicle-control treated *stk32a +/−* fish (blue). This phenotype is not enhanced in *stk32a −/−* fish treated with neomycin (magenta). Lines and shading represent mean ± SEM. Box plots show locomotor activity and sleep at night. Black bars and dark shading indicate night. n = number of fish. ns, P>0.05; **P<0.01; ***P<0.001; ****P<0.0001 by Mann-Whitney test **(e,f)** or Kruskal-Wallis test with Dunn’s test **(g,h)**.

In mice, *Stk32a* is expressed in the paraventricular nucleus of the hypothalamus, the homolog of the zebrafish NPO ^25^, and the suprachiasmatic nucleus **(Extended Data Fig. 6b,c)**. *Stk32a* is also expressed in cranial motor nuclei, including the trigeminal, facial, hypoglossal, and oculomotor nuclei **(Extended Data Fig. 6d-g)**, and in alpha motor neurons in the spinal cord ^26^. *Stk32a* is also expressed in sensory systems including the nucleus of the solitary tract **(Extended Data Fig. 6a)**, a brainstem population that transmits visceral sensory information to the brain ^27^, and in hair cells of the inner ear ^28^. The expression pattern of *Stk32a* is thus similar in mice and zebrafish.

### Phosphorylation of neurofilament proteins is reduced in *stk32a* mutants

To identify potential Stk32a substrates, we conducted phosphoproteomics, comparing adult zebrafish brains isolated from *stk32a −/−* and *stk32a +/+* siblings at night, when the *stk32a −/−* sleep phenotype is strongest. This analysis identified a small number of phosphopeptides that are less abundant in *stk32a −/−* fish **(Fig. 3a, Extended Data Fig. 7a, and Supplementary Table 1)**, most of which were from Nefma and neurofilament medium chain b (Nefmb), the zebrafish paralogs of the human NEFM protein. Several phosphopeptides from neurofilament light chain a (Nefla) and Neflb were also less abundant in *stk32a −/−* brains **(Fig. 3a and Extended Data Fig. 7a)**. Phosphorylation of these neurofilament subunits contributes to the assembly, turnover, and organization of neurofilaments that maintain the structure and function of large-caliber axons ^29^. Notably, reduced Nefma and Nefmb phosphorylation was primarily observed in the C-terminal region of these proteins **(Extended Data Fig. 7a)**, where phosphorylation provides resistance against proteolysis ^30^ and affects neurofilament assembly and transport ^31^. *nefma, nefmb, nefla,* and *neflb* are co-expressed with *stk32a* in CRNs, Mauthner neurons, and a subset of spinal cord motor neurons **(Fig. 2j and Extended Data Fig. 5d-i)**, which regulate locomotion ^22,23^. RNA-seq and scRNA-seq studies have also detected *nefma, nefmb, nefla,* and *neflb* transcripts in neuromast hair cells ^32–34^, which co-express *stk32a* **(Fig. 2d-e)**. These four neurofilament genes are also expressed in the anterior and posterior lateral line ganglia **(Fig. 3b and Extended Data Fig. 5d,f,h)** that innervate *stk32a*-expressing neuromasts **(Extended Data Fig. 8g)** to relay sensory information to the brain, as well as in reticulospinal neurons (RSNs) that regulate locomotion, and cranial nerve nuclei that have sensory and motor functions **(Extended Data Fig. 5d,f,h)** ^35^. Gene set enrichment analysis (GSEA) confirmed that significantly down-regulated phosphopeptides in *stk32a −/−* brains were enriched for genes associated with intermediate filament organization and intermediate filament-based processes **(Extended Data Fig. 7b)**. Human STK32A preferentially phosphorylates serine or threonine with acidic residues at positions P-4, P+1, P+2, and P+3 ^36^. Consistent with these findings, we observed that Nefma and Nefmb are phosphorylated at serine or threonine residues that have acidic residues at those positions **(Extended Data Fig. 7c,d)**. These results suggest that Stk32a phosphorylates neurofilament proteins in both sensory and motor systems.

### Ablation of neurofilament-expressing neurons results in increased sleep

A human GWAS reported that a SNP (rs17052966) near *NEFM* is associated with insomnia ^37^, suggesting that NEFM may affect human sleep, and supporting a link between phosphorylation of neurofilament proteins by Stk32a and sleep. To test the hypothesis that *NEFM* regulates sleep, we mutated the two zebrafish *NEFM* paralogs (*nefma* and *nefmb*), and found that *nefma* and *nefmb* single mutants (not shown) and double mutants (**Extended Data Fig. 8a,b**) lack locomotor activity or sleep phenotypes. This may result from the redundant action of other neurofilament proteins or from compensation for the chronic loss of *nefma* and *nefmb* during development. Since *nefma* and *nefmb* are largely co-expressed (**Fig. 3b**), we tested the effect of ablating *nefma*- expressing neurons using *Tg(nefma:Gal4);Tg(UAS:eNTR-mYFP)* fish ^38,39^. These fish exhibited normal locomotor activity and sleep compared to their non-transgenic siblings in the absence of metronidazole (MTZ) treatment **(Extended Data Fig. 8c,d)**, indicating that the transgenes themselves do not affect sleep. However, ablation of *nefma*-expressing neurons by treating these fish with MTZ **(Fig. 3c,d)** resulted in more sleep during the day and night due to more sleep bouts during the day and longer sleep bouts at night compared to identically treated non-transgenic siblings **(Fig. 3f and Extended Data Fig. 8e,f)**, similar to *stk32a* mutants **(Fig. 1c-e)**. Importantly, ablating these neurons had no significant effect on locomotor activity or waking activity during the day **(Fig. 3e and Extended Data Fig. 8e)**, suggesting the sleep phenotype is not due to abnormal locomotion or sickness. These results demonstrate that neurofilament-expressing neurons are required for normal sleep levels. However, since neurofilament genes are expressed in both motor and sensory systems, these experiments cannot determine which of these populations regulate sleep.

### Ablation of lateral line neuromasts phenocopies *stk32a* mutation

It was recently suggested that evolutionary loss of sleep in Mexican cave fish results from an increase in the number of lateral line neuromasts, which may result in increased arousal due to greater sensitivity to environmental stimuli ^40^. Since *stk32a* and neurofilaments are co-expressed in neuromast hair cells, and neurofilaments are also expressed in lateral line afferent neurons that transmit sensory information from neuromasts to the brain **(Extended Data Fig. 8g)**, we hypothesized that the *stk32a* mutant sleep phenotype is due, at least in part, to defective hair cell function. To test this hypothesis, we treated *stk32a +/−* and *stk32a −/−* fish with neomycin, which ablates neuromast hair cells ^41^ **(Extended Data Fig. 8h)**. Consistent with our hypothesis, neuromast-ablated *stk32a* +/− fish slept more than vehicle-control treated *stk32a +/−* fish **(Fig. 3h)** due to more daytime sleep bouts and longer nighttime sleep bouts **(Extended Data Fig. 8i,j)**, similar to *stk32a* −/− fish (**Fig. 1c-e**) and to fish whose *nefma*-expressing neurons are ablated (**Fig. 3f and Extended Data Fig. 8e,f**). Neomycin treatment did not further increase sleep in *stk32a −/−* fish **(Fig. 3h and Extended Data Fig. 8i,j)**, suggesting that neuromasts and *stk32a* affect sleep via the same pathway. These results suggest that *stk32a* mutants sleep more, at least in part, due to defective neuromast function, and thus a reduced ability to detect environmental stimuli, similar to the suppression of sensory neurons during invertebrate sleep ^42^.

### The serotonergic raphe promotes sleep via *stk32a*

To identify pathways through which *stk32a* regulates sleep, we tested for interactions between *stk32a* and known sleep regulators. We found that neither arousal induced by overexpression of *hypocretin (hcrt)* ^9^ **(Extended Data Fig. 9a)** nor sleep induced by overexpression of *npvf* ^17^ **(Extended Data Fig. 9b)**, loss of noradrenaline synthesis in *dopamine beta-hydroxylase* (*dbh)* mutants ^10^ **(Extended Data Fig. 9c)**, or administration of the α1 adrenergic receptor antagonist prazosin ^10^ **(Extended Data Fig. 9d)**, or melatonin ^11^ **(Extended Data Fig. 9e)** or were blocked in *stk32a −/−* fish. However, we did identify interactions between *stk32a*, the serotonergic (5-HT) raphe, and *nts*.

We previously showed that stimulation of 5-HT raphe neurons results in increased sleep, similar to *stk32a* mutation, whereas ablation of 5-HT raphe neurons reduces sleep, in both zebrafish and mice ^12^. These observations are consistent with the hypothesis that 5-HT raphe neurons promote sleep by inhibiting functions that require *stk32a*. To test this hypothesis, we first ablated the 5-HT raphe by MTZ treatment of *Tg(tph2:eNTR-mYFP)* fish. This resulted in more locomotor activity and less sleep in *stk32a* +/− fish, but this phenotype was abolished in *stk32a* −/− fish **(Fig. 4a,b)**. We next optogenetically stimulated 5-HT raphe neurons using *Tg(tph2:ChR2)* fish, which resulted in less locomotor activity and more sleep than non-transgenic siblings in *stk32a* +/− fish **(Fig. 4c,d)**. However, stimulation of 5-HT raphe neurons did not further increase sleep in *stk32a* −/− fish. Together, these experiments suggest that the raphe regulates sleep in a *stk32a*-dependent manner, possibly by inhibiting *stk32a-*expressing cells, or by inhibiting neurons that transmit signals from *stk32a*-expressing hair cells.

**Figure 4.**
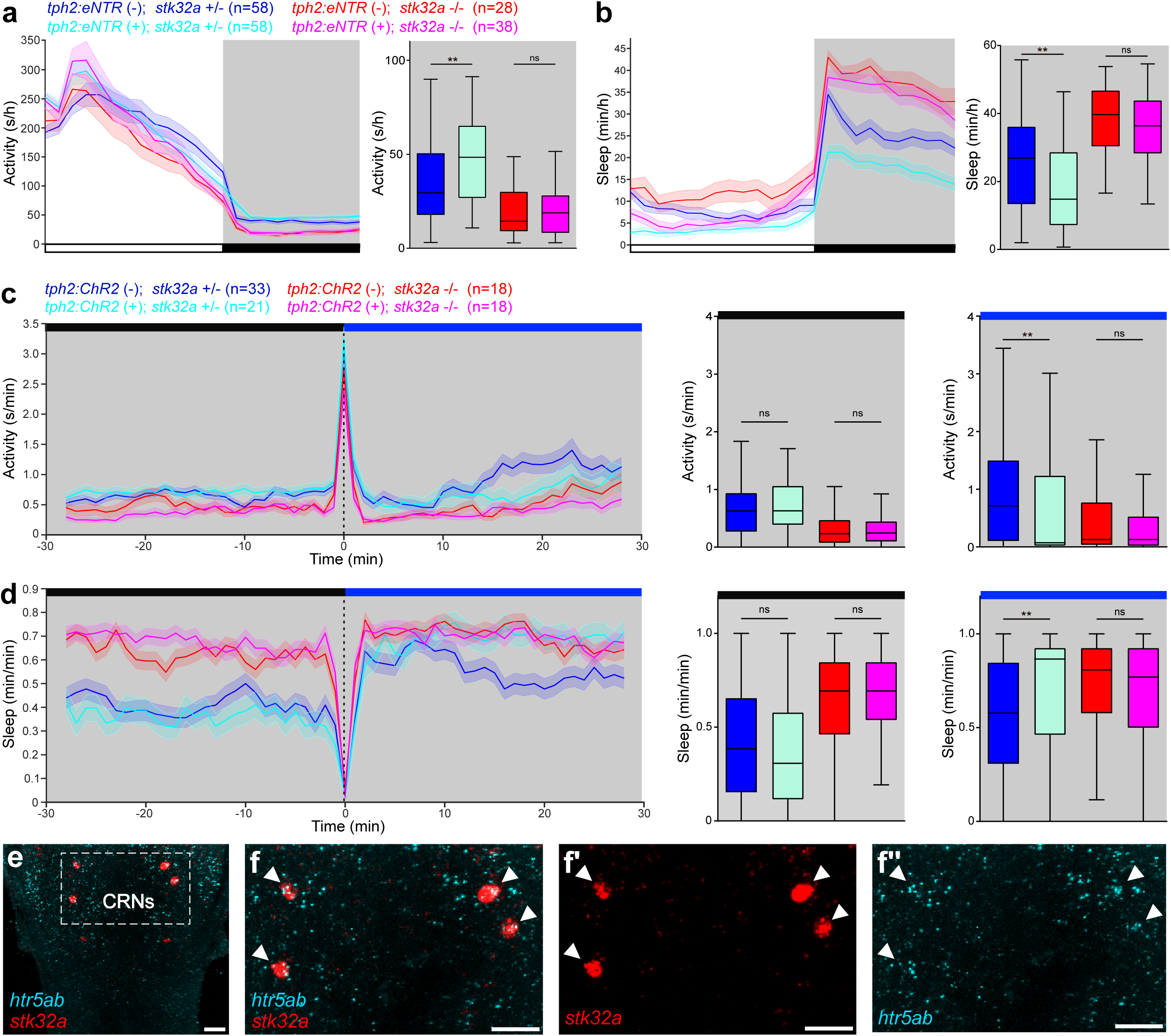
*stk32a* regulates sleep downstream of the 5-HT raphe. **(a,b)** Locomotor activity and sleep of 5-dpf *Tg(tph2:eNTR-mYFP)* (cyan and magenta) and non-transgenic sibling controls (blue and red) in the *stk32a +/−* or *stk32a −/−* mutant background following treatment with 5 mM MTZ. Ablation of the raphe increases activity and decreases sleep compared to non-transgenic controls in *stk32a +/−* fish, but not in *stk32a −/−* siblings. **(c,d)** Locomotor activity and sleep of 5- dpf *Tg(tph2:ChR2)* (cyan and magenta) and non-transgenic sibling controls (blue and red) before and during exposure to blue light in the *stk32a +/−* or *stk32a −/−* mutant background. Stimulation of the raphe decreases locomotor activity and increases sleep compared to non-transgenic controls in *stk32a +/−* fish, but does not further increase sleep in *stk32a −/−* siblings. Line graphs show mean ± SEM. Box plots quantify locomotor activity and sleep before and during blue light exposure. ns, P>0.05; **P<0.01 by Kruskal-Wallis test with Dunn’s test. **(e)** HCR of 6-dpf zebrafish shows that *htr5ab* and *stk32a* are co-expressed in CRNs. The boxed region in **(e)** is magnified in **(f-f”)**. Arrowheads indicate CRNs. Scale bars: 50 μm.

Next, we examined whether any 5-HT receptors are co-expressed with *stk32a* by performing HCR for excitatory (*htr2cl1, htr3a,* and *htr7c*) and inhibitory (*htr1aa, htr1ab, htr1b, htr1d,* and *htr5ab*) 5- HT receptors. We found that the inhibitory 5-HT receptor *htr5ab* is expressed in *stk32a*-expressing CRNs **(Fig. 4e,f)**. We did not detect 5-HT receptor expression in lateral line neurons or neuromasts, but these cells might express 5-HT receptors at levels too low to detect by HCR, or express a receptor that we did not test. Indeed, RNA-seq has detected expression of inhibitory 5- HT receptors *htr1aa, htr1ab, htr1b, htr1d, htr1e, htr1fa, htr5aa*, and *htr5ab* in neuromasts ^33,34,43^. These observations are consistent with the hypothesis that 5-HT can directly inhibit *stk32a*- expressing cells in sensory and motor systems.

### Neurotensin signaling promotes sleep and requires *stk32a*

We found that treating WT fish with an Nts receptor 1 (Ntsr1) agonist (PD149163) ^44^ resulted in less locomotor activity and more sleep due to more daytime sleep bouts and longer nighttime sleep bouts **(Extended Data Fig. 10a-d)**, similar to *stk32a* mutants. Conversely, treatment with an Ntsr1 antagonist (SR48692) ^45^ resulted in more locomotor activity and less sleep due to fewer and shorter daytime sleep bouts and shorter nighttime sleep bouts **(Extended Data Fig. 10e-h)**. These results suggest that Nts signaling promotes sleep in zebrafish, similar to mice ^46–48^, but in contrast to a prior study describing a wake-promoting role for Nts in zebrafish ^49^. Next, we asked whether Nts signaling promotes sleep via *stk32a*. Consistent with this hypothesis, *stk32a* −/− fish treated with the Ntsr1 agonist did not sleep more than *stk32a* −/− fish treated with DMSO vehicle control **(Fig. 5a,b)**, and *stk32a* −/− fish treated with the Ntsr1 antagonist did not sleep less than *stk32a* −/− fish treated with DMSO vehicle control **(Fig. 5c,d)**. These results suggest that Nts signaling acts upstream of Stk32a to promote sleep.

**Figure 5.**
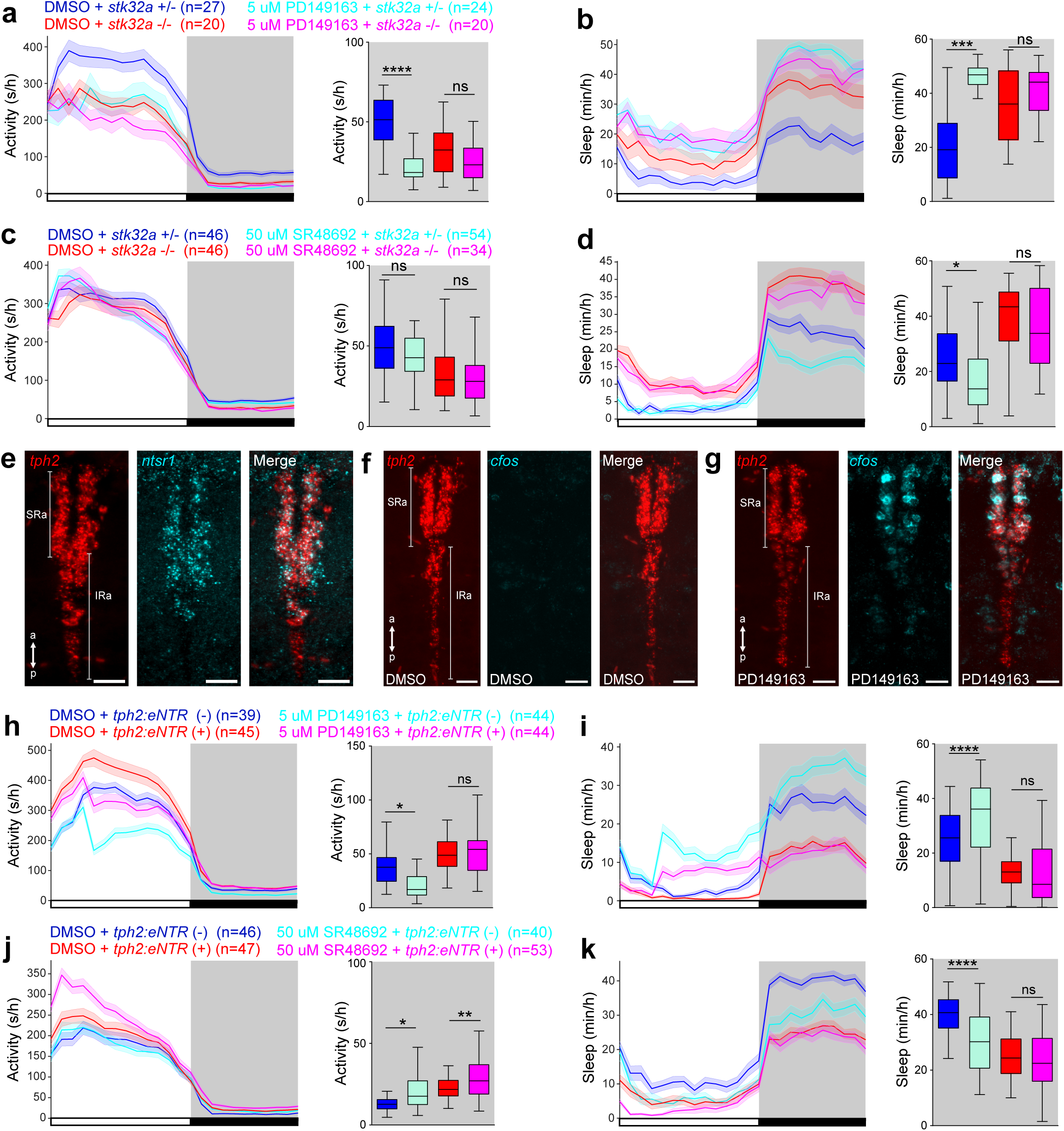
Ntsr1 signaling promotes sleep upstream of *stk32a* by activating the raphe. **(a,b)** Both *stk32a +/−* fish treated with 5 μM of Ntsr1 agonist PD149163 (cyan) and *stk32a −/−* fish treated with DMSO vehicle control (red) are less active and sleep more at night compared to DMSO treated *stk32a +/−* fish (blue). This phenotype is not enhanced in *stk32a −/−* fish treated with PD149163 (magenta). **(c,d)** *stk32a +/−* fish treated with 50 μM of Ntsr1 antagonist SR48692 (cyan) sleep less at night compared to both *stk32a +/−* (blue) and *stk32a −/−* fish (red) treated with DMSO vehicle control, but this phenotype is suppressed in *stk32a −/−* fish treated with SR48692 (magenta). **(e)** HCR reveals that *ntsr1* is expressed in the superior raphe (SRa) and rostral inferior raphe (IRa), labeled by *tph2* expression. **(f,g)** Treatment with 5 μM of Ntsr1 agonist PD149163, but not with DMSO vehicle control, results in *c-fos* expression in the SRa and rostral IRa. Scale bars: 25 μm. a, anterior; p, posterior. **(h,i)** Non-transgenic zebrafish treated with 5 μM of Ntsr1 agonist PD149163 (cyan) are less active and sleep more at night compared to raphe ablated *Tg(tph2:eNTR-mYFP)* zebrafish (red) and non-transgenic siblings treated with DMSO vehicle control (blue), but this phenotype is suppressed in raphe ablated *Tg(tph2:eNTR-mYFP)* zebrafish (magenta). **(j,k)** Non-transgenic zebrafish treated with 50 μM of Ntsr1 antagonist SR48692 (cyan) and raphe ablated *Tg(tph2:eNTR-mYFP)* fish (red) are more active and sleep less than non-transgenic WT siblings (blue) treated with DMSO vehicle control, but SR48692 treatment does not further decrease sleep in raphe-ablated *Tg(tph2:eNTR-mYFP)* fish (magenta). Fish of all genotypes were treated with MTZ in **(h-k)**. Lines and shading represent mean ± SEM. Box plots quantify locomotor activity and sleep at night. Black bars and dark shading indicate night. n = number of fish. ns, P>0.05; *P<0.05; **P<0.01; ***P<0.001; ****P<0.0001 by Kruskal-Wallis test with Dunn’s test.

### Neurotensin signaling promotes sleep by activating the 5-HT raphe

Since Nts signaling promotes sleep via Stk32a, we next asked if Nts signaling might act directly on *stk32a*-expressing neurons by performing HCR for *stk32a* and *ntsr1*. We did not detect any cells that co-express *stk32a* and *ntsr1* (data not shown). However, we found that *ntsr1* is co-expressed with *tph2*, a marker for 5-HT raphe neurons, throughout the superior raphe and in the rostral half of the inferior raphe **(Fig. 5e and Extended Data Fig. 10i,j)**. In addition, we found that administration of the Ntsr1 agonist PD149163, which promotes sleep **(Extended Data Fig. 10b)**, induced expression of *c-fos* throughout the superior raphe and in the rostral half of the inferior raphe **(Fig. 5f,g and Extended Data Fig. 10k,l)**, consistent with the expression pattern of *ntsr1*. This observation suggests that Nts signaling promotes sleep by stimulating 5-HT raphe neurons. If this hypothesis is correct, then ablation of 5-HT raphe neurons using MTZ treatment of *Tg(tph2:eNTR-mYFP)* fish should abolish the sedating effect of the Ntsr1 agonist, and should not enhance the arousing effect of the Ntsr1 antagonist. We found both of these predictions to be true. First, treatment with the Ntsr1 agonist increased sleep in non-transgenic control fish but not in raphe-ablated fish **(Fig. 5h,i)**. Second, while both ablation of the raphe and treatment of non-transgenic fish with the Ntsr1 antagonist reduced sleep, treating raphe-ablated fish with the Ntsr1 antagonist did not further decrease sleep compared to either single treatment **(Fig. 5j,k)**.

### Neurotensin signaling suppresses motor systems and sensory input from the lateral line

The above results suggest that Ntsr1 signaling promotes sleep by activating the 5-HT raphe, which in turn may inhibit sensory and motor systems that require Stk32a for normal function. To test the latter hypothesis, we performed calcium imaging using *Tg(nefma:Gal4); Tg(UAS:jGCaMP8f)* fish **(Fig. 6a and Extended Data Fig. 11c)** following treatment with DMSO vehicle or the Ntsr1 agonist PD149163, both at resting state and in response to a 600 Hz acoustic stimulus that was delivered every 2 minutes. We made three important observations. First, treatment with PD149163 resulted in a large reduction of resting jGCaMP8f fluorescence in sensory systems, including the anterior and posterior lateral line ganglia (ALLg, PLLg) **(Fig. 6b,c,f and Extended Data Fig. 11a)** and their afferent projections to the lateralis central lintel (LCL) **(Extended Data Fig. 11a)**, as well as in vestibulospinal neurons (VSNs) and the statoacoustic ganglia (SAG) **(Extended Data Fig. 11a).** Resting jGCaMP8f fluorescence was also reduced in motor systems, including the MNs and CRNs **(Fig. 6b,d-f and Extended Data Fig. 11a)**, as well as the oculomotor nucleus (nIII) and RSNs **(Extended Data Fig. 11a)**. Fluorescence did not decrease in the trochlear nucleus (nIV) or facial motor nucleus (nVII) **(Extended Data Fig. 11a)**, indicating that PD149163 treatment did not inhibit all neurons. Second, analysis of calcium transients (dF/F) (i.e. both spontaneous and stimulus-evoked responses) revealed stimulus-evoked jGCaMP8f responses in sensory (ALLg, PLLg, LCL, VSN, SAG) and some motor populations (MN and RSN), but not in CRNs or any cranial motor nuclei **(Fig. 6b-e,h and Extended Data Fig. 11h)**. Third, PD149163 treatment suppressed both spontaneous and stimulus-induced calcium transients in the sensory ALLg, PLLg, and LCL, but not in other sensory systems, or in any of the motor populations **(Fig. 6b-e,g,h and Extended Data Fig. 11b,d-h)**. These results suggest that activation of Ntsr1 signaling suppresses tonic firing of many sensory and motor populations, and also inhibits responses to stimuli in ALLg and PLLg neurons, which transmit sensory information from neuromasts to the brain. Taken together, our results suggest that Nts signaling promotes sleep by activating 5-HT raphe neurons, which in turn promote sleep by reducing tonic firing of several sensory and motor systems, and by inhibiting responses of the lateral line system to environmental stimuli **(Fig. 6i)**. This demonstrates that neuropeptidergic and neuromodulatory systems promote sleep homeostasis by inhibiting both sensory and motor systems, and establishes a mechanism through which increased homeostatic sleep pressure is coupled to increased arousal threshold, which is essential for sleep.

**Figure 6.**
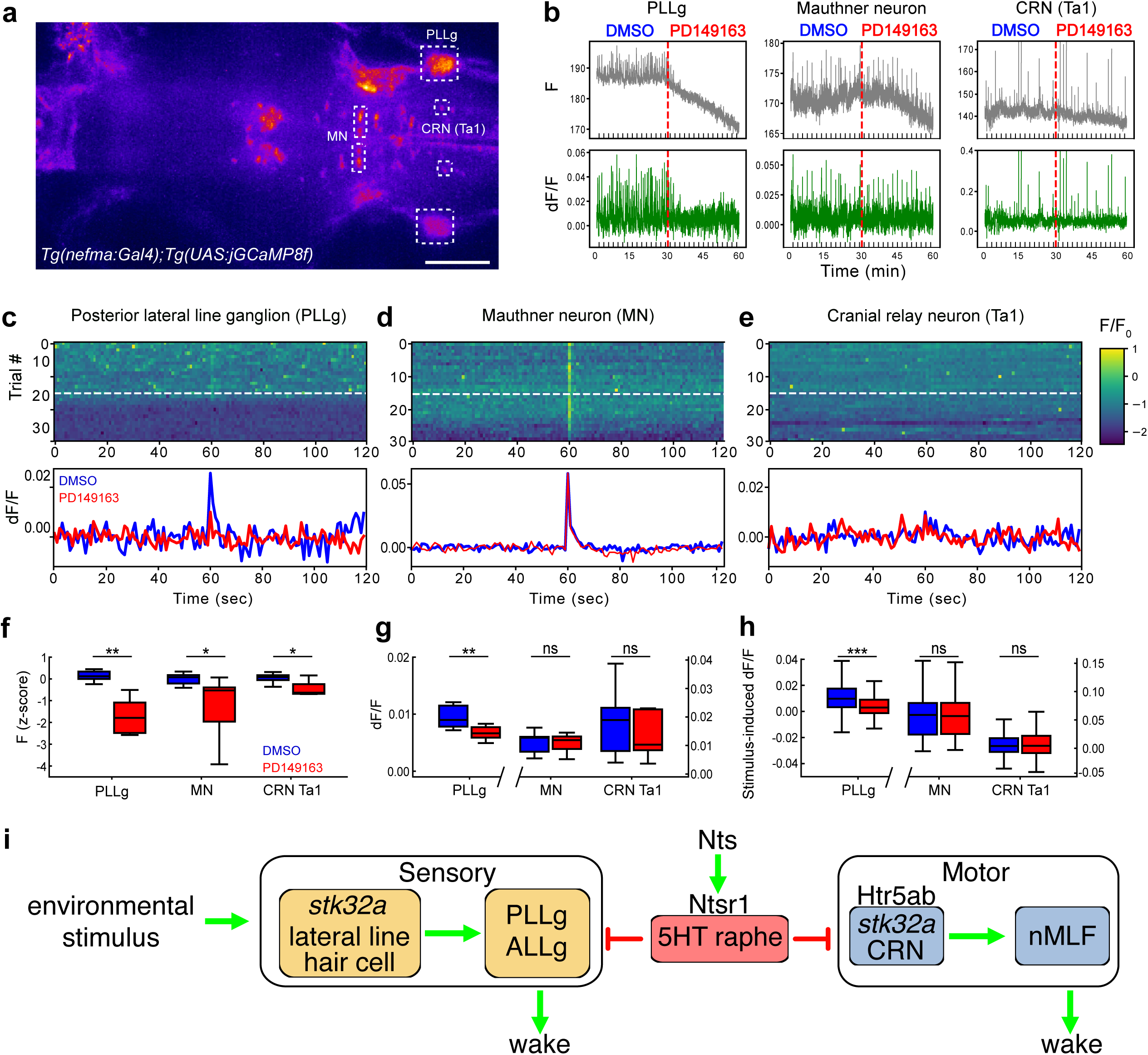
Nts signaling differentially suppresses neuronal activity in specific sensory and motor populations. **(a)** Maximum intensity projection of a 6-dpf *Tg(nefma:Gal4); Tg(UAS:jGCaMP8f)* fish. **(b)** Representative traces of raw fluorescence and dF/F from a fish treated with DMSO for 30 minutes and 5 μM of Ntsr1 agonist PD149163 for the next 30 minutes for the regions indicated in **(a)**. Red dashed lines indicate the start of PD149163 treatment. Black tick marks indicate acoustic stimulus presentations. **(c-e)** Heatmaps (top) and traces (bottom) of jGCaMP8f fluorescence normalized to the first frame (F/F0) or baseline (dF/F) for a fish subjected to a 600 Hz acoustic stimulus every 2 minutes during treatment with DMSO or PD149163 for the PLLg **(c)**, MN **(d)** and CRN (Ta1) **(e)**. White dashed lines in heat maps indicate the start of drug treatment. **(f-h)** Box plots comparing raw fluorescence **(f)**, dF/F **(g)**, and stimulus-induced responses **(h)** for DMSO and PD149163 treated fish. n = 7 fish. ns, P>0.05; *P<0.05; **P<0.01; ***P<0.001 by Mann-Whitney test. **(i)** Proposed model. Environmental stimuli promote wakefulness by stimulating lateral line hair cells, which relay this information to the brain via the PLLg and ALLg. This results in locomotor activity that is mediated by CRNs, which promote locomotion via the nMLF. The 5-HT raphe promotes sleep homeostasis by inhibiting both sensory (PLLg and ALLg) and motor (CRN) populations, possibly in part by binding of 5-HT to the inhibitory Htr5ab on CRNs. Nts promotes sleep by stimulating 5-HT raphe neurons, likely via Ntsr1 that is expressed in 5-HT raphe neurons. *stk32a* is required in lateral line hair cells and/or CRNs for homeostatic regulation of sleep by the Nts-raphe system.

## DISCUSSION

Recent studies have identified several protein kinases that regulate sleep, including the sleep-promoting SIK1, SIK2, SIK3, ERK, and CaMKII, and the wake-promoting protein kinase A ^50^, which are broadly expressed and likely regulate sleep via brain-wide effects. Indeed, sleep and wake states are correlated with hypophosphorylation and hyperphosphorylation, respectively, of many synaptic proteins, which may serve as a molecular basis for sleep homeostasis ^50^. In contrast, here we identify *stk32a* as a novel protein kinase with a critical role in sleep whose expression is restricted to specific sensory, neuropeptidergic, and motor populations, suggesting it regulates sleep via specific neural circuits. Indeed, we found that *stk32a* functions downstream of Nts signaling and the 5-HT raphe to promote wakefulness. Consistent with its restricted expression pattern, loss of *stk32a* affects the phosphorylation of far fewer proteins than broadly expressed kinases like *Sik3* ^51,52^. Since *stk32a* mutants show increased sleep, one might expect that Stk32a normally phosphorylates proteins in wake-promoting neurons. Indeed, most of the hypophosphorylated proteins in *stk32a* mutants are neurofilament proteins, which are specifically expressed in RSNs, CRNs, lateral line ganglion neurons, and neuromast hair cells that we found are required for normal levels of wakefulness. The identification of Nefma and Nefmb as Stk32a phosphorylation targets that are relevant to sleep is supported by a GWAS that identified SNPs near *NEFM* that are associated with insomnia ^37^. Neurofilaments are important for the structure and function of neurons with large caliber axons, such as RSNs, CRNs and lateral line ganglion neurons, and may serve a similar function in hair cells. Phosphorylation of neurofilament subunits modulates their assembly, turnover, and organization within the axonal cytoskeleton ^53^, but we did not observe obvious axonal or neurofilament defects in *stk32a −/−* or *nefma −/−; nefmb −/−* zebrafish (data not shown), similar to a prior study of *NEFM* mutant mice ^54^. Additional studies are needed to determine how reduced neurofilament phosphorylation in *stk32a* mutants and loss of neurofilament genes affects neurons and hair cells at the molecular and ultrastructural levels, and how these changes affect sleep.

The two-process model, proposed over 40 years ago ^1^, postulates that sleep is regulated by a homeostatic process that increases with time spent awake and decreases during sleep, and a circadian process that determines sleep timing. Sleep is also rapidly affected by light and dark, a phenomenon known as masking ^55^. We found that *stk32a* is dispensable for circadian- and light-dependent regulation of sleep, but acts downstream of the raphe to regulate sleep homeostasis, and that Nts signaling promotes sleep upstream of both the raphe and *stk32a*. We previously showed that raphe plays a key role in sleep homeostasis ^12^, in part in response to hypothalamic *npvf*-expressing neurons that activate the rostral half of the inferior raphe ^17,56^. Here we show that Nts signaling promotes sleep, likely by activating the superior raphe and the rostral half of the inferior raphe, suggesting that the raphe integrates multiple sleep-promoting signals to control homeostatic sleep drive.

How might homeostatic sleep drive be implemented in the brain? Our experiments suggest this involves inhibition of specific sensory and motor systems that express *stk32a*. First, *stk32a* is expressed in hair cells of lateral line neuromasts ^32–34^ and the ear, consistent with a role for *stk32a* in sensing acoustic and water flow stimuli. We observed that Nts signaling activates the raphe and suppresses both resting and stimulus-induced activity of lateral line neurons that transmit arousing sensory information from hair cells to the brain, thus linking the sleep homeostat to the increased arousal threshold that is essential for sleep. *stk32a* is also expressed in the olfactory epithelium and retina, suggesting the Nts-raphe-Stk32a pathway may also promote sleep drive by suppressing responses to olfactory and visual stimuli. Second, *stk32a* is expressed in CRNs and Mauthner neurons that promote motor activity via nMLF neurons ^23^. We found that sleep induced by Nts signaling, and thus likely by the 5-HT raphe, suppresses resting, but not stimulus-evoked, activity of these neurons and other RSNs, likely via the inhibitory Htr5ab receptor in CRNs and other inhibitory 5-HT receptors that are expressed in RSNs ^57^. Together, these observations suggest that the raphe integrates neuropeptidergic signals that promote homeostatic sleep pressure, and then drives sleep by suppressing both sensory and motor systems. Our findings are consistent with the hypothesis that a function of the raphe is to integrate information over time to drive behavior ^58–60^. Moreover, the coordinated inhibition of both sensory and motor systems by the raphe links the classical two-process model, which postulates a role for homeostatic sleep pressure in promoting sleep ^1^, with the more recently proposed “arousal-action” model, which postulates that sleep involves direct inhibition of motor systems ^61^. Our findings thus reveal an evolutionarily conserved role for *stk32a* that links the two-process and arousal-action models to the suppression of sensory and motor systems that is essential for sleep.

## METHODS

### EXPERIMENTAL MODEL AND SUBJECT DETAILS

#### Zebrafish husbandry

All zebrafish experiments were performed in accordance with the Institutional Animal Care and Use Committee (IACUC) guidelines and by the Office of Laboratory Animal Resources at the California Institute of Technology (animal protocol 1836). Zebrafish of the TLAB strain were generated by mating TL and AB fish strains obtained from the Zebrafish International Resource Center (ZIRC, Oregon). TLAB (WT) animals were used to generate mutant and transgenic animals. Adult zebrafish were maintained on a 14:10 hour light:dark cycle with lights on at 9 a.m. and off at 11 p.m. All experiments used healthy and experimentally naïve zebrafish larvae between 4-7 dpf, a developmental stage prior to sexual differentiation. Zebrafish larvae were raised in petri dishes, with 50 animals per plate, containing E3 medium (5 mM NaCl, 0.16 mM KCl, 0.33 mM CaCl2, 0.33 mM MgSO4). *Tg(nefma:Gal4)* transgenic zebrafish were a gift from Shin-ichi Higashijima ^39^. This line contains a *hsp70l:LOXP-RFP-LOXP-GAL4* transgene inserted into the *nefma* genomic locus. We used fish in which the *LOXP-RFP-LOXP* sequence was removed by mating to zebrafish that ubiquitously express Cre recombinase, resulting in fish that express Gal4 in *nefma*-expressing cells.

#### Zebrafish mutagenesis

Zebrafish mutants were generated using CRISPR/Cas9 by injecting a single guide RNA (sgRNA) (Integrated DNA Technologies) targeting genes of interest and Cas9 protein (Alt-R^TM^ S.p. Cas9 Nuclease, Cat No 1081058, Integrated DNA Technologies) into TLAB embryos at the single cell stage. sgRNA sequences were selected using CHOPCHOP V2 ^62^. T7 endonuclease (Cat No. M0689, New England Biolabs) was used to confirm the cutting efficiency of sgRNAs. The nature of each mutation was determined by Sanger sequencing. For *stk32a* mutants, the sgRNA sequence 5’-ATTTTGAGAGCCATCGGCAA-3’ introduced a 20 bp deletion (5’- CAAAGGCAGTTTTGGGAAGG-3’) at the end of exon 2 of *stk32a* (ENSDARG00000096879), causing a frameshift of the open reading frame, and introduction of a premature stop codon, resulting in a predicted 45 amino acid truncated mutant protein compared to the 403 amino acid WT protein (see Fig. S1B). This mutant protein lacks the kinase domain and is likely to be a complete loss of function allele. To generate *nefma* and *nefmb* mutants, two sgRNAs flanking the coding region of each gene were co-injected with Cas9 protein in zebrafish embryos at the single cell stage. Flanking PCR primers designed to amplify genomic sequence only if the sequence between both sgRNAs was excised were used to identify mutants, and Sanger sequencing was used to confirm the nature of each deletion. The excised regions did not include any other annotated genes. The *nefma* (ENSDARG00000021351) mutant was generated using two sgRNAs with target sequences 5’-ACACATCACGAGTGCTTGGG-3’ and 5’- AGTAATATACAATCGAATGG-3’, which induced a 4650 bp deletion that includes 252 bp of the 5’UTR, the entire open reading frame, and 408 bp of the 3’UTR. The *nefmb* (ENSDARG00000043697) mutant was generated using two sgRNAs with target sequences 5’- GTCTTCATCGATTTTCCGAG-3’ and 5’-GAACAAACCCCCAACCCGAG-3’, which induced a 3072 bp deletion that includes the 90 bp of sequence upstream of the 5’UTR, the 5’UTR, the open reading frame, and 87 bp of the 3’ UTR. To genotype fish following experiments, genomic DNA was prepared using standard methods ^63^, and PCR was performed using the primers described below, with PCR products run on a 4% agarose gel. *stk32a* mutants were genotyped using the primers 5’-TTGTGTTTTGTCTCTCCACAGTC-3’ and 5’-TCATGTAGGTTATAAATACAAAGCAGA-3’, which produced 110 bp and 90 bp bands for the WT and mutant alleles, respectively. *nefma* mutants were genotyped using the primers 5’- TGTATATCGCACACTTGATTAGCC-3’, 5’-GCTACTTCGTCAGAAGAACACG-3’, and 5’- GACCATTGATCCGTCGATGT-3’, which produced 1 band for homozygous WT (118 bp), 2 bands for heterozygous mutants (118 and 101 bp), and 1 band for homozygous mutants (101 bp). *nefmb* mutants were genotyped using the primers 5’-GCATTGAGGTGAATTGCTGA-3’, 5’- GCGTTCTGTCCCGTTGTATT-3’, and 5’- TCACCACCAAAACCACACAT-3’, which produced 1 band for WT (135 bp), 2 bands for heterozygous mutants (135 and 122 bp), and 1 band for homozygous mutants (122 bp).

#### Zebrafish transgenesis

We generated *Tg(5xUAS:mYFP-T2A-eNTR)* transgenic fish by cloning a mYFP-T2A-eNTR open reading frame ^64,65^ downstream of a 5xUAS element and upstream of a beta-globin intron, followed by an SV40 polyA sequence, into a plasmid containing NRSE elements ^66,67^ and flanking Tol2 transposase recognition sequences ^68^. Stable transgenic lines were generated by injecting the plasmid and *tol2 transposase* mRNA into embryos at the single cell stage. Transgenic animals were identified by mYFP expression. For simplicity, *Tg(5xUAS:mYFP-T2A-eNTR)* fish are referred to as *Tg(UAS:eNTR)* in the figures.

#### Mouse husbandry: pharmacology experiments

All experimental procedures followed National Institutes of Health guidelines and were approved by the Animal Care and Use Committee at the Institute of Neuroscience, Chinese Academy of Sciences. WT mice (C57BL/6) were purchased from institute-approved vendors (Shanghai LingChang Experiment Animal Co., China). Male mice (8 weeks at the time of surgery) were used. Mice were housed in rooms (temperature: 23 ± 1°C; humidity: 50–70%) under a 12:12 hour light:dark cycle (light on from 7 a.m. to 7 p.m.) with ad libitum access to food and water.

#### Mouse husbandry: CRISPR/Cas9 and HCR experiments

All procedures were approved by the Institutional Animal Care and Use Committee of the National Institute of Biological Sciences, Beijing (NIBS). Rosa26-Cas9 mice (026179, JAX) were obtained from the Jackson Laboratory. All mice were provided with food and water ad libitum and housed under controlled conditions: temperature (23 ± 1°C), relative humidity (50% ± 10%), and a 12:12- hour light:dark cycle (light on from 9 a.m. to 9 p.m.).

### METHOD DETAILS: ZEBRAFISH

#### Pharmacology

We treated larval zebrafish with TAE684 (4 µM, Selleckchem, S1108), SB-245391 (4 µM, Structural Genomics Consortium, Internal ID UNC-ZDG-5-27), SR 48692 (50 µM, Sigma, SML0278), PD 149163 tetrahydrochloride hydrate (5 µM, Sigma, PZ0175), neomycin sulfate (200 µM, SigmaMillipore, PHR1491), melatonin (5 µM, Sigma, M5250), DASPEI (0.005%, SigmaMillipore, D0815), and prazosin hydrochloride (2 µM, Sigma, P7791). All drugs were dissolved in a final concentration of 0.2% DMSO (VWR, MK494802), which does not itself affect behavior. Drugs were added directly to each well of a 96-well plate containing E3 medium and 4- dpf fish. Dose-response experiments were performed to identify drug concentrations that induced robust behavioral phenotypes, without apparent toxicity or abnormal responses to gentle stimuli. For sleep experiments, drugs were added during the evening at 4-dpf and behavioral data was reported for the fifth day and night of development. For neomycin experiments, fish were treated with neomycin for 1 hour in the morning of 4-dpf and washed with E3, followed by five hours of recovery before being placed in a 96-well plate

#### Sleep behavioral experiments

Sleep behavioral experiments were performed using a videotracking system as previously described ^9^. Zebrafish were raised in petri dishes (50 per dish) on a 14:10 hour light:dark (LD) cycle at 28.5°C with lights on a 9 a.m. and off at 11 p.m. For optogenetic experiments, animals were raised in dim white light to prevent stimulation of ChR2 during development. Starting at 4- dpf, individual fish were placed in each well of a 96-well plate (Whatman, 7701-1651), filled with 650 µL E3 embryo medium. Locomotor activity was quantified using an automated videotracking system (Viewpoint Life Sciences) with a Dinion one-third Monochrome camera (Dragonfly 2, Point Grey) fitted with a variable-focus megapixel lens (Computar, M5018-MP) and an infrared filter. The movement of each fish was recorded using quantization mode. The 96-well plate and camera were housed inside a custom Zebrabox (Viewpoint Life Sciences), with white light illumination from 9 a.m. to 11 p.m., and continuous infrared light illumination for video recording. The 96-well plate was housed in a chamber filled with recirculating water to maintain a constant temperature of 28.5°C. The parameters used for detection (detection threshold: 15, burst: 29, freeze: 3, bin size: 60 s) were empirically determined. For most experiments, behavior was monitored for 24 hours, during the 5th day and night of development. For free-running behavioral experiments, fish were raised and tested in normal LD cycles until the 5th night of development, and then were shifted to either constant light or constant dark and monitored for an additional 48 hours. Peak to trough ratios were calculated by dividing average 1 hour locomotor activity during the middle of subjective day 7 (ZT55) by average 1 hour locomotor activity during the middle of subjective night 7 (ZT67) for constant light and constant dark experiments. To analyze behavior in the absence of entrained circadian rhythms, animals were raised in constant light and behaviorally tested in constant light. Fish were placed into the videotracker at 4-dpf and data was reported for 48 hours corresponding to the 5th and 6th day and night of development. Data were analyzed using custom MATLAB (Mathworks) scripts ^69^. Zebrafish were genotyped by PCR after each experiment to identify mutant, transgenic, and WT siblings.

#### Arousal threshold assay

The arousal threshold assay was performed as previously described ^10^ using a videotracking system with an Arduino-based automated driver to control two solenoids (Guardian Electric, 28P- I-12) that delivered a tap at 14 different intensities to a 96-well plate containing zebrafish larvae. Individual 4-dpf fish were placed into each well of a 96-well plate filled with E3 medium. On the 5th night of development, a series of taps were applied to the plate in pseudo-random intensities from 12:30 a.m. to 7:30 a.m. with an inter-trial interval (ITI) of 1 minute. Previous studies showed that an ITI of 15 seconds is sufficient to prevent behavioral habituation ^70^. The background probability of movement was calculated by identifying the fraction of fish for each genotype that moved 5 seconds prior to all stimuli delivered during the experiment (14 tap intensities x 30 trials per experiment = 420 data points per fish; average background movement). This value was subtracted from the average response fraction value for each tap event (corrected response = average response – average background movement). Behavioral responses to the stimuli were monitored using the videotracking software and analyzed using MATLAB (Mathworks) and Excel (Microsoft). Statistical analysis was performed using the Variable Slope log(dose) response fitting module of Prism (GraphPad). The ratio of fish that responded to each tap was plotted against the tap intensity, and the EP50 (effective power 50) was calculated as the tap intensity that resulted in a half-maximal response.

#### Sleep deprivation

Zebrafish were placed in the videotracker at 4-dpf and were behaviorally monitored starting at 5- dpf. Sleep deprivation was performed during the first 6 hours of the 6th night of development by maintaining daytime levels of white light illumination as previously described ^12^. Animals that failed to show at least a 50% reduction in sleep during the first 4 h of sleep deprivation (SD Sleep) compared to sleep during the first 4 h of the previous night (Baseline Sleep) were excluded from the analysis. For each genotype, accumulated recovery sleep was calculated by subtracting Baseline Sleep (first 4 hours of night 5 after lights off) from cumulative Recovery Sleep (first 4 hours of night 6 after sleep deprivation and lights off) at each corresponding time point with the data binned in 20-minute intervals. Only 4 hours of Recovery Sleep (and hence Baseline Sleep) were used to ensure that all sleep data were confined to the circadian night.

#### Optogenetic stimulation

The videotracking system was modified to include an array of blue LEDs (470 nm, Luxeon V-star, MR-BO0040-10S) mounted 15 cm above and 7 cm away from the center of the 96-well plate to ensure uniform illumination, as previously described ^10^. The LEDs were controlled using a custom driver and software written in BASIC stamp editor. A power meter (Laser-check, 1098293) was used before each experiment to confirm light intensity (∼500 μW at the surface of the 96-well plate). In the afternoon at 4-dpf, zebrafish were placed in each well of a 96-well plate in a videotracker, and allowed to habituate overnight with exposure to a single 30 minute blue light stimulus at 12 a.m. On the 5th night of development, 6 trials were performed in which fish were exposed to blue light for 30 minutes, with 60 minutes of recovery in between each trial, starting at 12 a.m. The behavior of each fish was monitored for 30 minutes before and after blue light onset. Since the onset of blue light induces a startle response and a short burst of locomotor activity, we excluded 5 minutes of data centered around the peak of the startle response, as well as 2 minutes before light offset, from the analysis. For baseline, we used a time period equal to blue light exposure, but prior to blue light onset.

#### Chemogenetic ablation

Fish were treated with 5 mM metronidazole (VWR, 443-48-1) diluted in E3 medium containing 0.2% DMSO from 2-4 dpf, refreshed every 24 hours, as previously described ^12^. Fish were kept in dim light during the day during the treatment period to prevent MTZ photodegradation. In the morning of 4-dpf, fish were rinsed three times with E3 medium and allowed to recover for at least 3 hours prior to being transferred to 96-well plates for sleep experiments. The reported data is from the 5th day and night of development.

#### Hybridization chain reaction (HCR)

HCR was performed as previously described ^71^. Whole zebrafish larvae were fixed in 4% PFA in PBS overnight with nutation at 4°C, and subsequently washed with 0.1% Tween-20/PBS (PBSTw). Samples were permeabilized with 100% methanol at −20°C for 10 minutes, rehydrated with graded dilutions of methanol/PBSTw, and washed with PBSTw. To improve visualization, samples were treated with a bleaching solution (1.5% hydrogen peroxide and 0.1% potassium hydroxide) for 20 minutes, and then washed with PBSTw. Samples were pre-hybridized for 60 minutes and then hybridized with HCR probes (2 pmol of each probe set in 500 μL) for 48 hours at 37°C with nutation. Samples were washed with a wash buffer (Molecular Instruments) at 37°C and equilibrated with 5x SSCT. Samples were then transferred to the amplification buffer with 30 pmol of each hairpin that were snap-cooled, and the reaction was carried out at room temperature in the dark overnight. Samples were then washed with 5xSSCT and gradually transferred to Vectashield antifade mounting medium (Vector Laboratories) in 5xSSCT prior to imaging using a Zeiss LSM 880 confocal microscope.

#### Phosphoproteomics

Quantitative phosphoproteomic experiments were performed on whole-brain lysates of 12-month-old adult zebrafish frozen in liquid nitrogen at ZT 17 (2 a.m.). Brains were quickly dissected from frozen adults and stored on dry ice. Brain samples were homogenized in lysis buffer (50 mM HEPES pH 7.4, 150 mM NaCl, 2.5% SDS, 2 mM MgCl2) supplemented with protease and phosphatase inhibitor cocktail tablets (Thermo Scientific, 78447). Homogenates were incubated at room temperature for 30 minutes and then centrifuged at 15,000 g for 20 minutes. The supernatant was recovered, and protein concentration was determined using the bicinchoninic acid assay ^72^ (Pierce, 23225). Samples were reduced, digested, subjected to C18 solid-phase extraction, and vacuum-centrifuged to dryness to obtain desalted peptides. Titansphere Phos-TiO MP Kit (GL Sciences Inc, 5010-21283) was used to capture and enrich for phosphopeptides. Peptide and phosphopeptide mixtures were labelled with tandem mass tag (TMT) prior to separation using liquid chromatography-mass spectrometry (LC-MS). Samples were analyzed using an Orbitrap Eclipse Tribrid mass spectrometer with FAIMS Pro coupled with an EASY-nLC 1200 liquid chromatography pump. We compared LC-MS/MS analysis with and without FAIMS and concluded that we can detect 15% more proteins with FAIMS, which is especially useful for low abundant proteins. Raw data was searched with Proteome Discoverer SEQUEST (version 2.5, Thermo Scientific) against the *Danio rerio* (zebrafish) FASTA file downloaded from UniProt. The post-analysis of TMT-labeling quantitative phosphopeptides was performed using Tidyproteomics 1.7.3 ^73^. Protein/peptide ratio calculation was performed using a pairwise ratio-based method. Statistical analysis was performed using two-sided moderated Student t-test as implemented in the limma package to determine statistical significance ^74^. Volcano and heatmap plots were generated using the ggplot2 package v.3.4.4 ^75^. Gene Ontology (GO) enrichment analysis was conducted using overrepresentation analysis (ORA) via the clusterProfiler R package ^76^. The mass spectrometry proteomics data have been deposited to the ProteomeXchange Consortium via the PRIDE partner repository with the dataset identifier PXD064245 ^77^.

#### Calcium imaging

Calcium imaging was performed using 6-dpf *Tg(nefma:Gal4); Tg(UAS:jGCaMP8f)* fish using a custom-built light-sheet microscope with a 20x water immersion objective (Thorlabs, N20X-PFH). Fish were anesthetized by immersion in 0.016% w/v tricaine dissolved in E3 medium and embedded in 1.5% low-melting agarose. Agarose was dissected away from the gills and mouth to facilitate drug perfusion into the fish. Images were acquired using a 488 nm laser at 1 volume/second, with 6 μm between sections, and 50 sections/brain volume (300 μm total volume). Laser power at the objective was quantified using a power meter (ThorLabs, PM121D). Acoustic stimuli were delivered from an 8 ohm 200 MW speaker embedded in the bottom of the imaging chamber. The stimulus consisted of a 10 millisecond 600 Hz pulse that was delivered every 2 minutes during the 90 minute experiment. E3 medium was perfused through the imaging chamber for 30 minutes, followed by 0.1% DMSO in E3 for 30 minutes, followed by 5 μM Ntsr1 agonist PD149168 (Sigma, PZ0175) for 30 minutes. All solutions were heated to 28.5°C. dF/F was calculated as previously described ^78^ by normalizing dF to the baseline, which was defined as the 20th percentile in a 5-minute window centered at each time point.

### METHOD DETAILS: MOUSE PHARMACOLOGY EXPERIMENTS

#### Surgical procedures

All surgical procedures were conducted under general anesthesia using continuous isoflurane (5% for induction; 1.5%–2% for maintenance). Depth of anesthesia was monitored continuously and adjusted when necessary. Following induction of anesthesia, mice were placed on a stereotaxic frame with a heating pad. To implant EEG and EMG electrodes, stainless steel screws for EEG were inserted into the skull above the visual cortex and above the frontal cortex, two insulated EMG electrodes were inserted into the neck muscle, and a reference electrode was attached to a screw inserted into the skull above the cerebellum.

#### Polysomnography recordings

Mice were connected with a flexible cable for EEG/EMG activity. The EEG and EMG signals (high-pass filtered at 0.5 Hz and digitized at 1 kHz) were acquired using an RHD acquisition board and RHX software (Version 3.03, Intan Technologies). Video recordings were started simultaneously to observe behavior. Mice were placed individually in rice buckets with bedding, food, and water three days in advance. Four rice buckets were placed in a sound-attenuation box (68 cm x 62 cm x 74 cm) under a 12:12-hr light:dark cycle (light on at 7 a.m.). After a three-day adaptation period, three days of baseline recording was started and the box was not opened during the entire recording.

#### Drug delivery

NVP-TAE 684 (HY-10192,MCE,40 mg/kg) was injected at 6 p.m. and recording started at 7 p.m. NVP-TAE 684 was dissolved in DMSO and then diluted in sterile saline to a final value of 10% DMSO. For the control group, 10% DMSO in sterile saline was injected. The experimental group and control group were placed in the same box and injection was randomized.

#### Analysis of sleep architecture

EEG spectral analysis was carried out using a fast Fourier transform with a frequency resolution of 0.18 Hz. Brain states were automatically scored every 5 seconds in MATLAB (Accusleep_GUI) and validated manually by trained experimenters. Brain state classification was performed according to established criteria ^79,80^: Wakefulness was defined as desynchronized EEG and high EMG activity; NREM sleep was defined as synchronized EEG with high-amplitude delta activity (0.5–4 Hz) and low EMG activity; REM sleep was defined as high power at theta frequencies (6– 10 Hz) and low EMG activity. To analyze changes in the sleep–wake cycle after NVP-TAE 684 injection, we used the 12-h polysomnography recording one hour after injection to calculate the time, bout number, and bout length of each brain state. Eight mice were injected with NVP-TAE 684 and seven mice were injected with vehicle control.

### METHOD DETAILS: MOUSE CRISPR/CAS9 AND HCR EXPERIMENTS

#### Plasmid Construction

Triple-target CRISPR/Cas9 constructs for ABC-KO were created by designing sets of three sgRNA sequences for *Stk32a*, *Stk32b*, and *Stk32c* using the CRISPR direct website (https://crispr.dbcls.jp). Annealed oligonucleotides for each sgRNA were cloned into pAAVU6- sgRNA-hSyn-Cre-2A-EGFP-KASH-WPRE (Addgene, 60229), followed by sequential subcloning to generate pAAV-3xsgRNA.

#### AAV Packaging and Purification

Recombinant AAVs (rAAVs) were produced by transfecting AAVpro 293T cells, confirmed mycoplasma-free, with specific plasmids using polyethylenimine MAX. Cells were co-transfected with PHP.eB, pHelper, and transfer plasmids. After 72 hours, cells were harvested, lysed through freeze-thaw cycles with vortexing, and treated with benzonase nuclease. The lysate was centrifuged, and the supernatant containing rAAVs was purified via iodixanol gradient centrifugation. Purified rAAVs were concentrated using Amicon Ultra filters and formulated in phosphate-buffered saline containing Pluronic F68. Final rAAV titers were determined by qPCR, with linearized AAV plasmids as standards.

#### EEG/EMG-Based Sleep Analysis

Surgeries for EEG/EMG electrode implantation were conducted on 11- to 13-week-old male mice. Mice were anesthetized with isoflurane (4% induction, 2% maintenance). The skull was cleaned with hydrogen peroxide to enhance bonding with dental cement. EEG electrode pins were implanted into the dura at coordinates (−1.27, 0, 0), (−1.27, 5.03, 0), (1.27, 5.03, 0), and (1.27, 0, 0), while EMG wires were inserted into the neck muscles. Both were secured using dental cement.

Mice were acclimated to the recording setup for one week before baseline sleep recordings (three days). After a 6-hour sleep deprivation period, recovery sleep was recorded for 18 hours. EEG data underwent fast Fourier transform (FFT) analysis (1–30 Hz, 1 Hz intervals), and 20-second epochs were categorized as NREMS, REMS, or Wake using semi-automated software. Epoch classifications were manually reviewed for accuracy. Sleep duration plots during NREMS were calculated and averaged over three days.

#### Sleep Deprivation

To prevent curling, mice were placed in shallow water (∼1 cm height) during a 6-hour sleep deprivation period at the beginning of the light phase, with electrodes protected by parafilm. After sleep deprivation, EEG/EMG recordings resumed for 18 hours (remaining 6 hours of the light phase and 12 hours of the dark phase). Accumulated recovery NREMS was calculated by subtracting pre-deprivation NREMS durations from post-deprivation values at corresponding time point.

### QUANTIFICATION AND STATISTICAL ANALYSIS

For all behavioral experiments (mouse and zebrafish) the unit of analysis for statistics is a single animal, except for optogenetic and calcium imaging experiments where the unit of analysis for statistics is a single trial. Line graphs represent mean ± SEM. Tukey box plots were used for data presentation. Data points outside the Tukey range were not shown to facilitate data presentation, but were included in statistical analyses. Statistical analyses were performed using Prism 10 (GraphPad). Shapiro-Wilk normality tests indicated that most behavioral data were not normally distributed. Therefore, we used non-parametric tests for statistical analyses (Mann-Whitney test for two unpaired groups, Kruskal-Wallis test with Dunn’s correction for multiple comparisons for more than two unpaired groups). For mouse genetic experiments, we used 2-way ANOVA, followed by Dunnett’s test to compare every mean to the control mean, or Sidak’s test to compare a set of means. Data are considered to be statistically significant if P < 0.05.

## ACKNOWLEDGEMENTS

We thank Viveca Sapin, Uyen Pham, Daisy Chilin, Axel Dominguez, Alex Mack, Caressa Wong, Barbara Orozco, and Stephanie Li for zebrafish husbandry assistance, Claire Wyart and Takashi Kawashima for sharing unpublished data, Shinichi Higashijima for sharing transgenic fish, Minoru Koyama for helpful discussions, and Catherine Oikonomou and Katie Kindt for comments on the manuscript. This work was supported by funding from the National Institutes of Health (D.A.P. R35 NS122172 and UF1 NS126562, D.A.P. and T.F.C. R03 TR003353), the Howard Hughes Medical Institute (S.N. and M.B.A.), and the New Cornerstone Science Foundation and National Major Project of China Science and Technology Innovation 2030 for Brain Science and Brain-Inspired Technology (Q.L. 2021ZD203400). S.T. was supported by a postdoctoral fellowship from the Natural Science and Engineering Research Council (NSERC), a Banting Postdoctoral Fellowship, and a Sleep Research Society Career Development Award.

## AUTHOR CONTRIBUTIONS

Conceptualization: ST, DAP

Methodology: ST, AA, GO, SN, BG, TC

Investigation: ST, JE, CZ, XL, ML, AA, BG, TS, CG, HH, MY, SL, FW, TYW

Visualization: ST, JE, CZ, XL, TYW, DAP

Funding acquisition: ST, MBA, TFC, MX, QL, DAP

Project administration: MBA, TFC, MX, QL, DAP

Supervision: MBA, TFC, MX, QL, DAP

Writing – original draft: ST, DAP

Writing – edit and revision: ST, JE, CZ, XL, GO, TYW, MBA, TFC, MX, QL, DAP

## MATERIALS AND CORRESPONDENCE

Correspondence and material requests should be addressed to David A. Prober.

## TABLE LEGENDS

**Supplementary Table 1. Phosphoproteomic analysis of *stk32a −/−* vs. *stk32a +/+* zebrafish brains.**

---

**Extended Data Figure 1.**
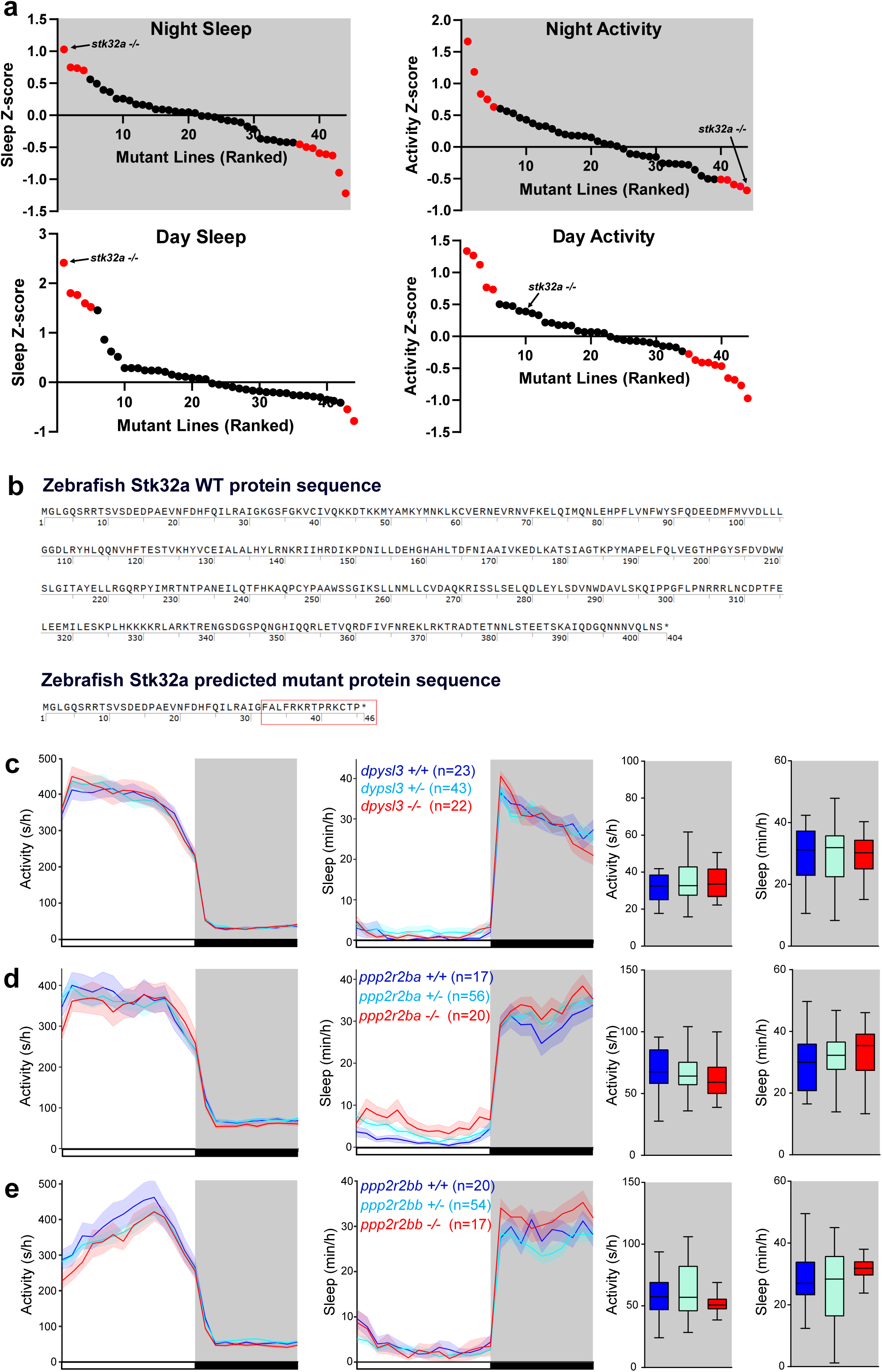
Zebrafish genetic screen summary. **(a)** Nighttime and daytime sleep and locomotor activity for each homozygous mutant in the genetic screen, normalized to their WT sibling controls and ranked by standard deviations from the mean (Z-score). Twenty mutants showed significant changes in daytime or nighttime sleep or locomotor activity (indicated in red). **(b)** Amino acid sequence of WT zebrafish Stk32a and the truncated mutant Stk32a protein. The red box indicates the new amino acid sequence introduced as a result of the frameshift mutation. **(c-e)** Locomotor activity and sleep for *dpysl3* **(c)**, *ppp2r2a* **(d)**, and *ppp2r2b* **(e)** mutants. Mean ± SEM (left) and box plots (right) are shown. Black bars and gray shading indicate night. n = number of fish. None of the mutants show statistically significant differences from sibling controls by Kruskal-Wallis test with Dunn’s test.

**Extended Data Figure 2.**
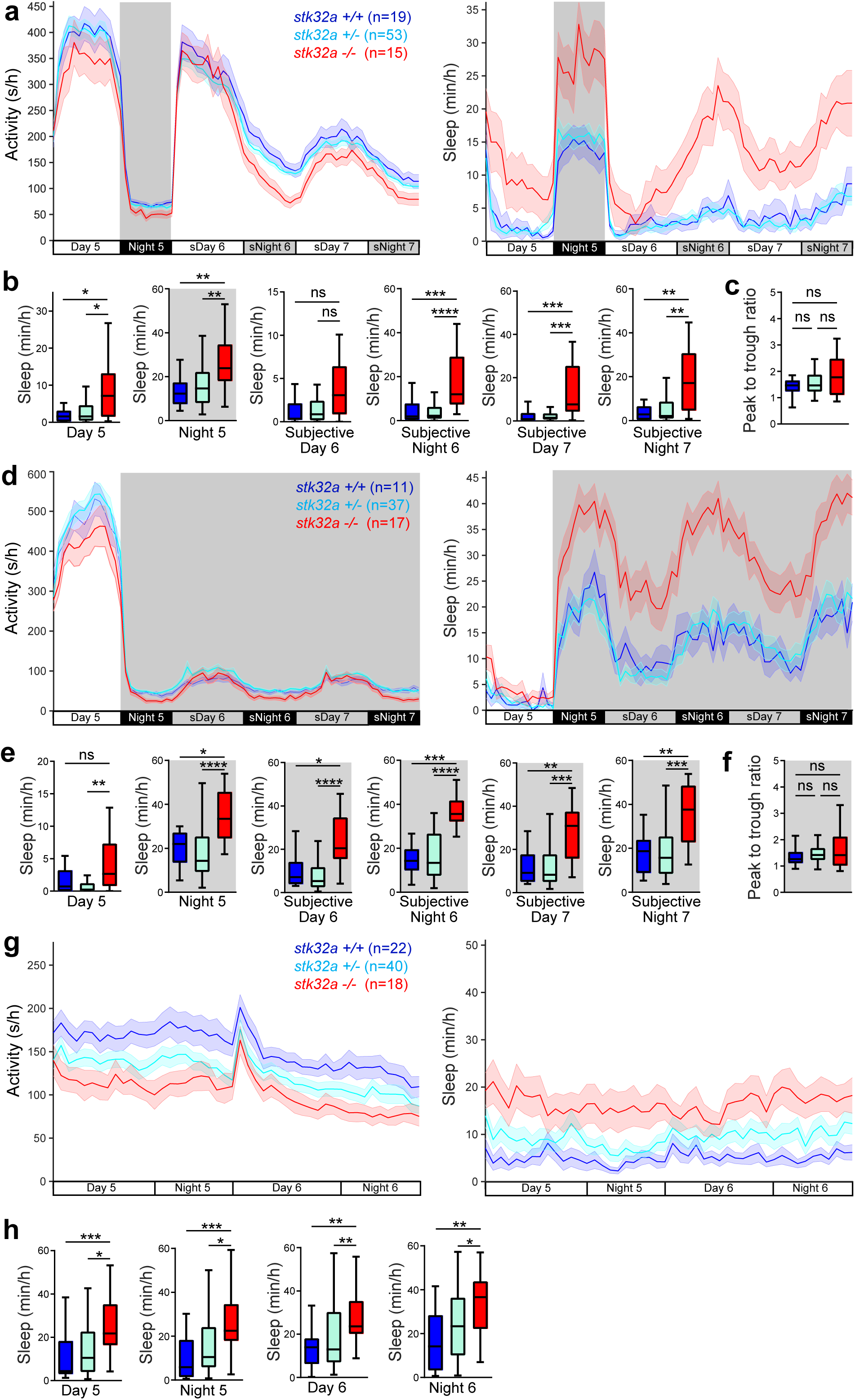
Zebrafish *stk32a* mutant phenotype is independent of lighting conditions and circadian rhythms. **(a,d)** Locomotor activity and sleep of *stk32a −/−* zebrafish and sibling controls raised on a 14:10 hour light:dark (LD) cycle and shifted to constant light **(a-c)** or constant dark **(d-f)** at 6-dpf. The *stk32a* mutant phenotype persists during both subjective day (sDay) and subjective night (sNight) in both conditions. Locomotor activity peak to trough ratios of subjective day 7 (average 1 hour activity at ZT55) to subjective night 7 (average 1 hour activity at ZT67) are shown for all genotypes in constant light **(c)** and constant dark **(f)**. Gray shading indicates darkness. Black bars indicate night and gray bars indicate subjective night **(a)** or subjective day **(d)**. **(g,h)** Locomotor activity and sleep of *stk32a −/−* zebrafish and sibling controls raised and tested in constant light. The transient increase in locomotor activity on the 6^th^ day of development is an artifact caused by addition of water to each well to offset evaporation during the experiment. Mean ± SEM and box plots are shown. n = number of fish. ns, P>0.05; *P<0.05; **P<0.01; ***P<0.001; ****P<0.0001 by Kruskal-Wallis test with Dunn’s test.

**Extended Data Figure 3.**
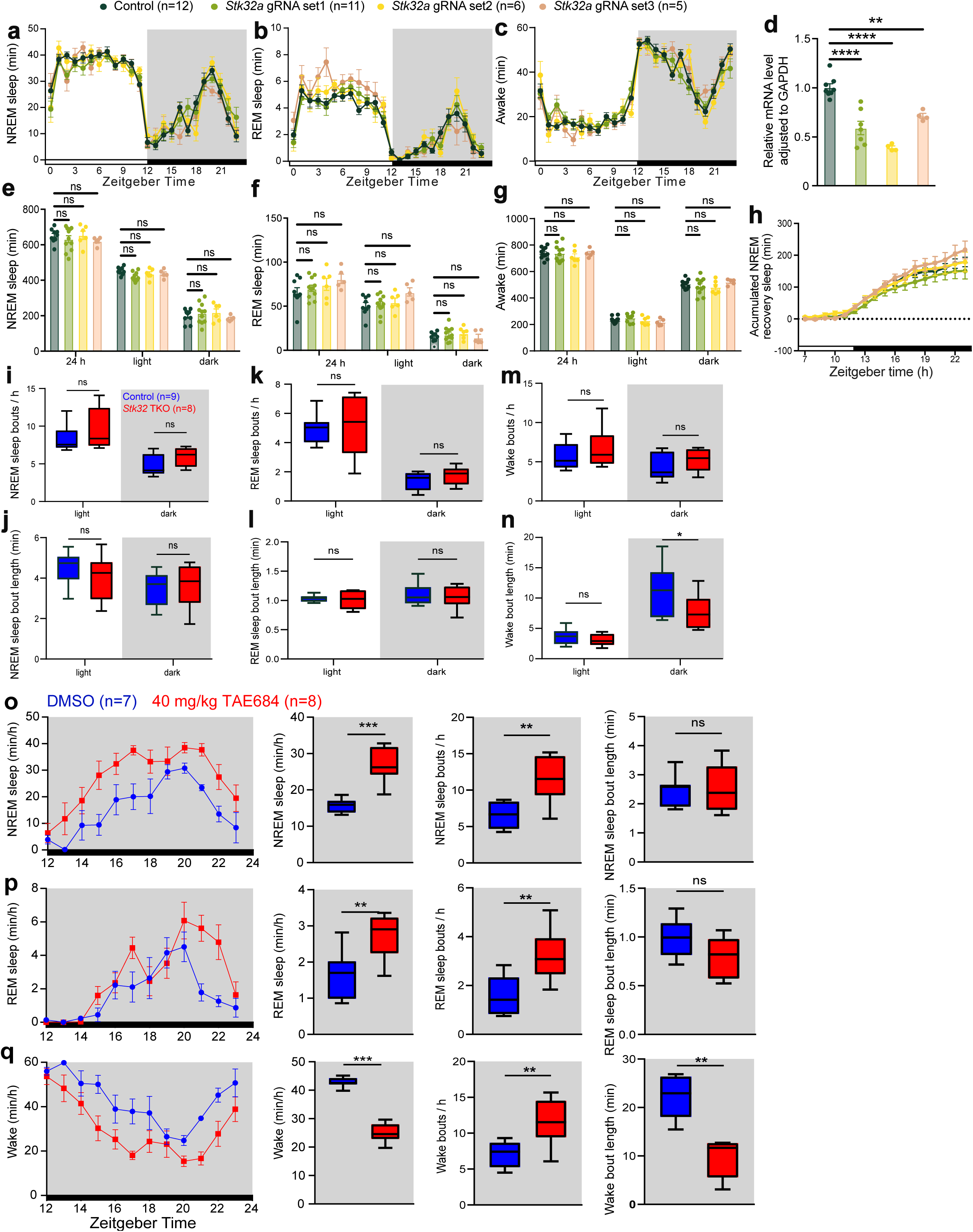
Genetic knock-out and pharmacological inhibition of Stk32 in mice. **(a-h)** Adult brain chimeric knockout (ABC-KO) of *Stk32a* in mice using different gRNA sets significantly reduced the expression of *Stk32a* compared to controls **(d)**, but did not significantly affect time in NREM sleep **(a,e)**, REM sleep **(b,f)**, or wake **(c,g)**, or accumulated NREM recovery sleep following sleep deprivation **(H)**, compared to controls. Mean ± SEM are shown. **(i-n)** Box plots quantify bout number (top) and bout length (bottom) for NREM sleep **(i,j),** REM sleep **(k,l)**, and wake **(m,n)** in *Stk32* TKO mice compared to controls. **(o-q)** Mice injected with 40 mg/kg TAE684 at zeitgeber time (ZT) 11 and monitored from ZT 12-23 spent more time in NREM and REM sleep and less time awake compared to DMSO vehicle control. Box plots quantify NREM sleep **(o)**, REM sleep **(p)**, and wake **(q)** time (left), bout number (middle), and bout length (right). Line graphs show mean ± SEM. Black bars and dark shading indicate night. n = number of mice. ns, P>0.05; *P<0.05; **P<0.01; ***P<0.001; ****P<0.0001 by two-way ANOVA with Dunnett’s test **(d-h)** or Mann-Whitney test **(i-q)**.

**Extended Data Figure 4.**
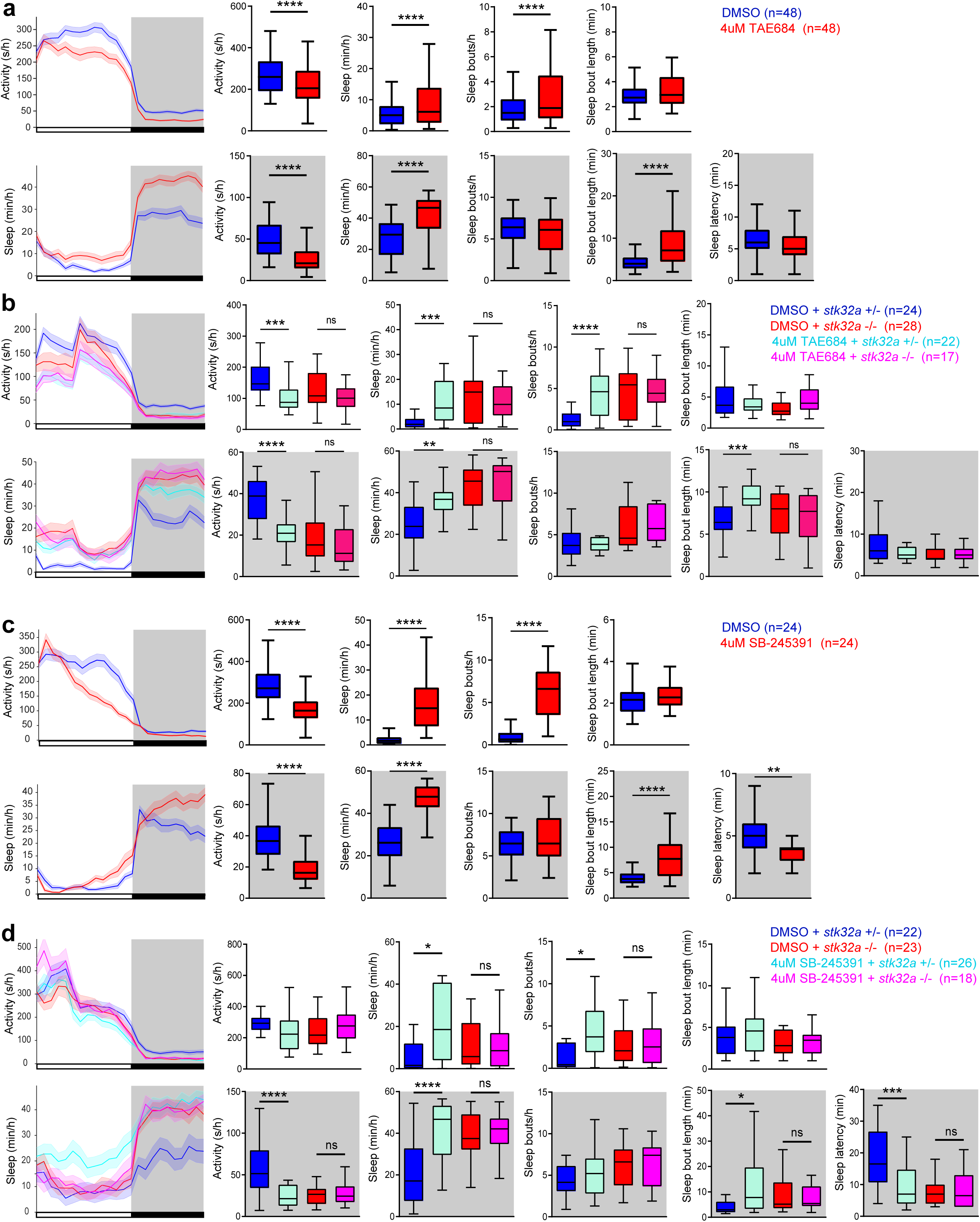
Effects of pharmacological inhibition of Stk32a on zebrafish sleep. **(a-d)**. Treating zebrafish with 4 μM of Stk32a inhibitor TAE684 or SB-245391 compared to DMSO vehicle control results in less locomotor activity and more sleep in WT fish **(a,c)** and in *stk32a* +/− fish **(b,d)**, but does not further decrease locomotor activity or increase sleep in *stk32a* −/− fish **(b,d)**. Box plots quantify locomotor activity, sleep, number of sleep bouts, and sleep bout length during the day and night, and sleep latency at night, in WT zebrafish treated with 4 μM TAE684 or DMSO **(a)**, *stk32a +/−* and *stk32a −/−* zebrafish treated with 4 μM TAE684 or DMSO **(b)**, WT zebrafish treated with 4 μM SB-245391 or DMSO **(c)**, and *stk32a +/−* and *stk32a −/−* zebrafish treated with 4 μM SB-245391 or DMSO **(d)**. Line graphs show mean ± SEM. Black bars and dark shading indicate night. n = number of fish. ns, P>0.05; *P<0.05; **P<0.01; ***P<0.001; ****P<0.0001 by Mann-Whitney test **(a,c)** or Kruskal-Wallis test with Dunn’s test **(b,d)**.

**Extended Data Figure 5.**
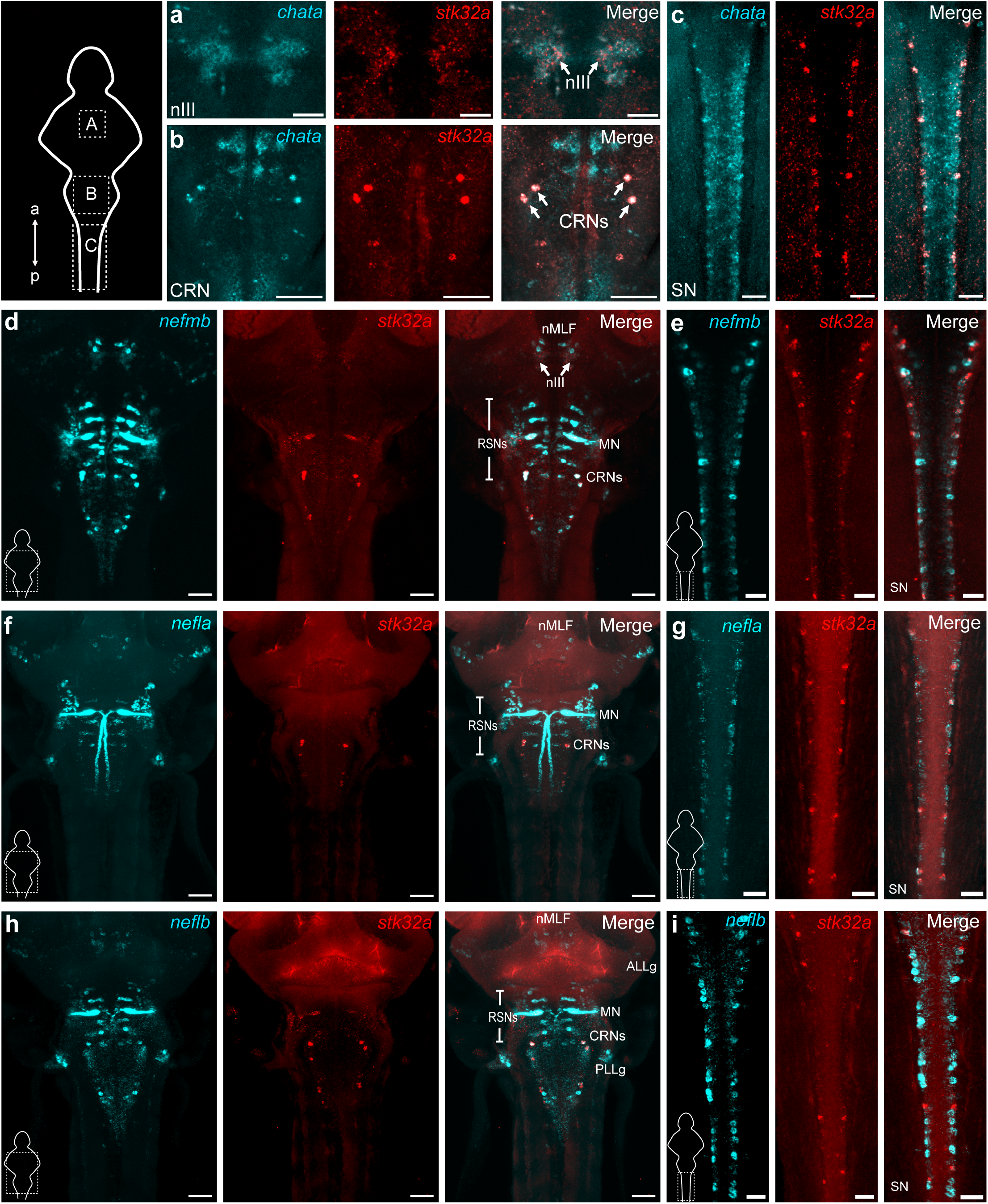
*stk32a* is co-expressed with neurofilament genes in zebrafish. HCR in 6-dpf zebrafish shows co-expression of *chata* **(a-c)**, *nefmb* **(d,e)***, nefla* **(f,g)**, and *neflb* **(h,i)** with *stk32a* in the oculomotor nucleus (nIII), Mauthner neurons (MN), cranial relay neurons (CRNs) and spinal cord neurons (SN). Scale bars: 50 μm.

**Extended Data Figure 6.**
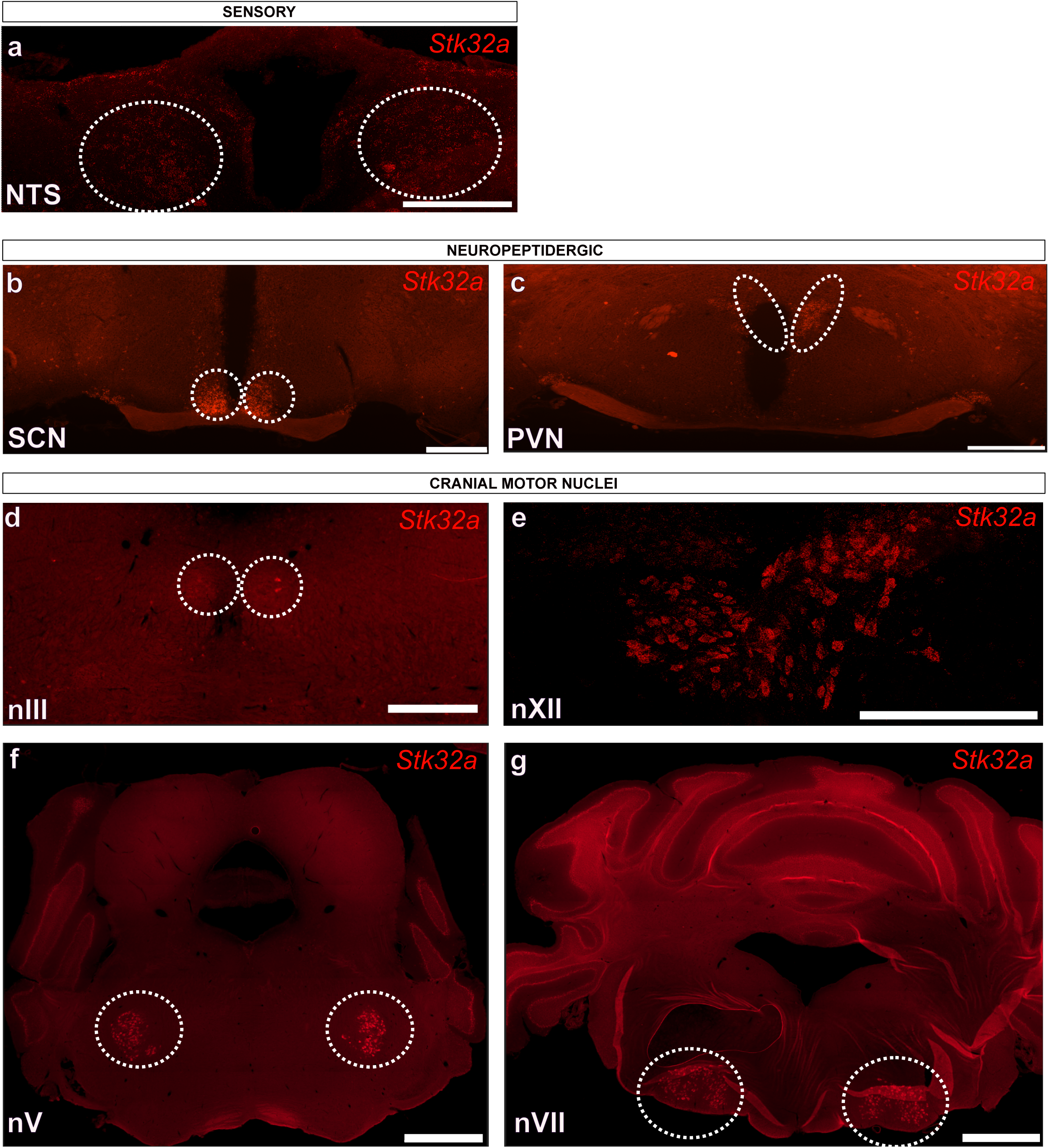
*Stk32a* is expressed in sensory, neuropeptidergic, and motor regions in mice. **(a)** RNAscope of adult mouse brain slices detected *Stk32a* mRNA in the nucleus of the solitary tract (NTS), the primary visceral sensory relay station in the brain. **(b,c)** *Stk32a* mRNA was also detected in neuropeptidergic cells of the hypothalamus, including the suprachiasmatic nucleus (SCN) **(b)** and paraventricular nucleus (PVN) **(c)**. **(d-g)** *Stk32a* was also detected in several cranial motor nuclei including the oculomotor nucleus (nIII) **(d)**, hypoglossal nucleus (nXII) **(e),** trigeminal nucleus (nV) **(f)**, and facial nucleus (nVII) **(g)**. Scale bars: 250 μm **(a)**, 500 μm **(b-e)**, 1000 μm **(f,g)**.

**Extended Data Figure 7.**
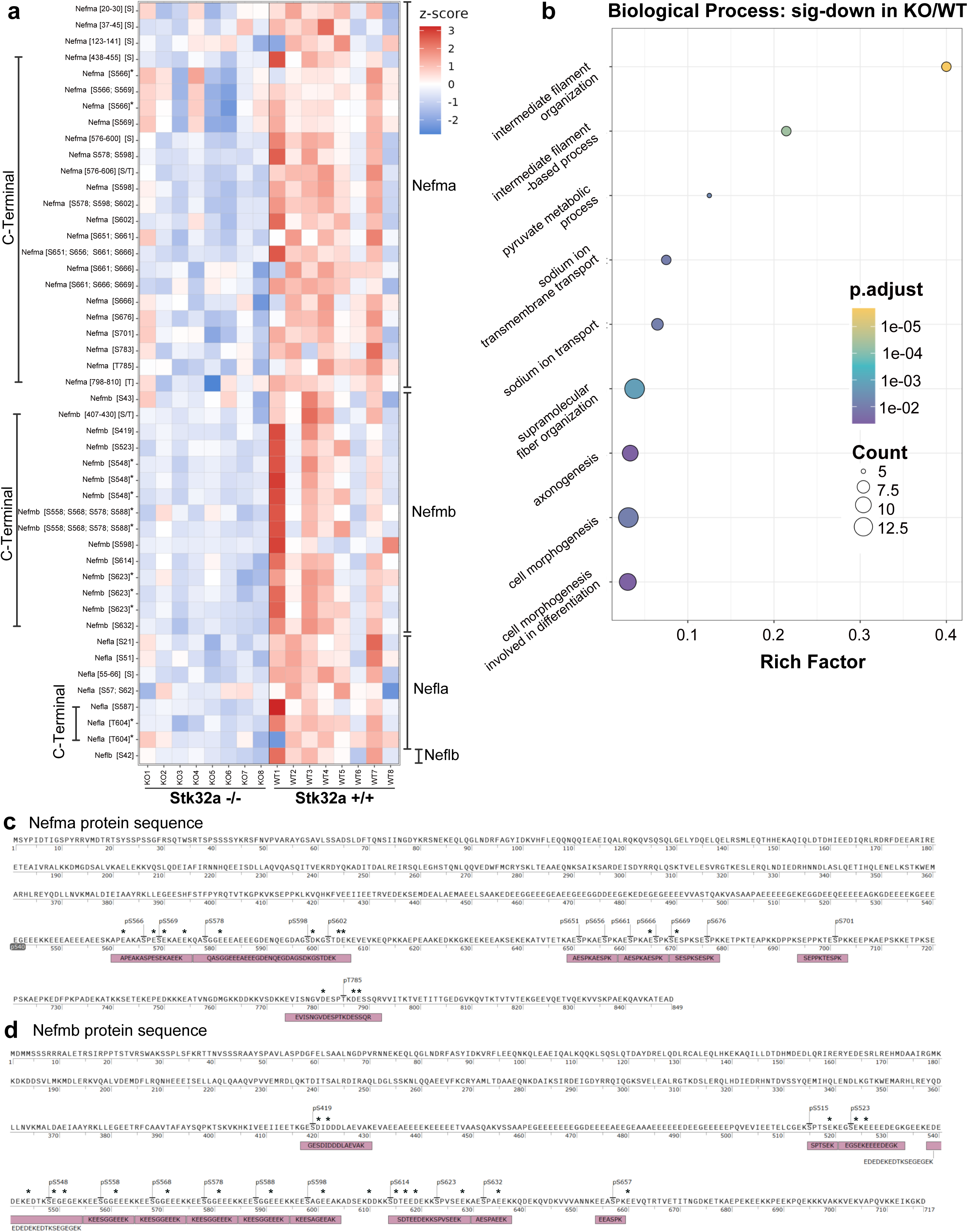
Reduced Nefma and Nefmb phosphopeptides in *stk32a* −/− zebrafish brains. **(a)** Heatmap comparing abundances of specific Nefma, Nefmb, Nefla, and Neflb phosphopeptides for individual *stk32a* −/− (KO) and *stk32a* +/+ (WT) brain samples. The phosphorylation site for each peptide is indicated. Undetermined phosphorylation sites are indicated as [S] or [T] after the detected amino acid sequence. Asterisks indicate repeated sites identified from missed cleavage peptides containing overlapping sequences with shared modified residues. **(b)** Gene set enrichment analysis (GSEA) of phosphopeptides significantly downregulated in *stk32a −/−* brains compared to *stk32a* +/+ controls reveals enrichment for Gene Ontology (GO) biological process terms, with intermediate filament organization and intermediate filament-based processes as the most enriched terms. The rich factor represents the ratio of differentially abundant proteins annotated in each GO biological process term to all differentially abundant proteins. The size of each point (count) corresponds to the number of differentially abundant proteins, and the color of each point represents the false-discovery-rate (FDR)-adjusted p-value (p. adjust). **(c,d)** The amino acid sequences of zebrafish Nefma **(c)** and Nefmb **(d)** are shown, including phosphopeptide sequences (magenta boxes) and phosphorylation sites (each annotated with its amino acid residue number) identified by phosphoproteomics. Asterisks indicate acidic amino acid residues (aspartate or glutamate) at positions −4, +1, +2, or +3 relative to a phosphorylation site.

**Extended Data Figure 8.**
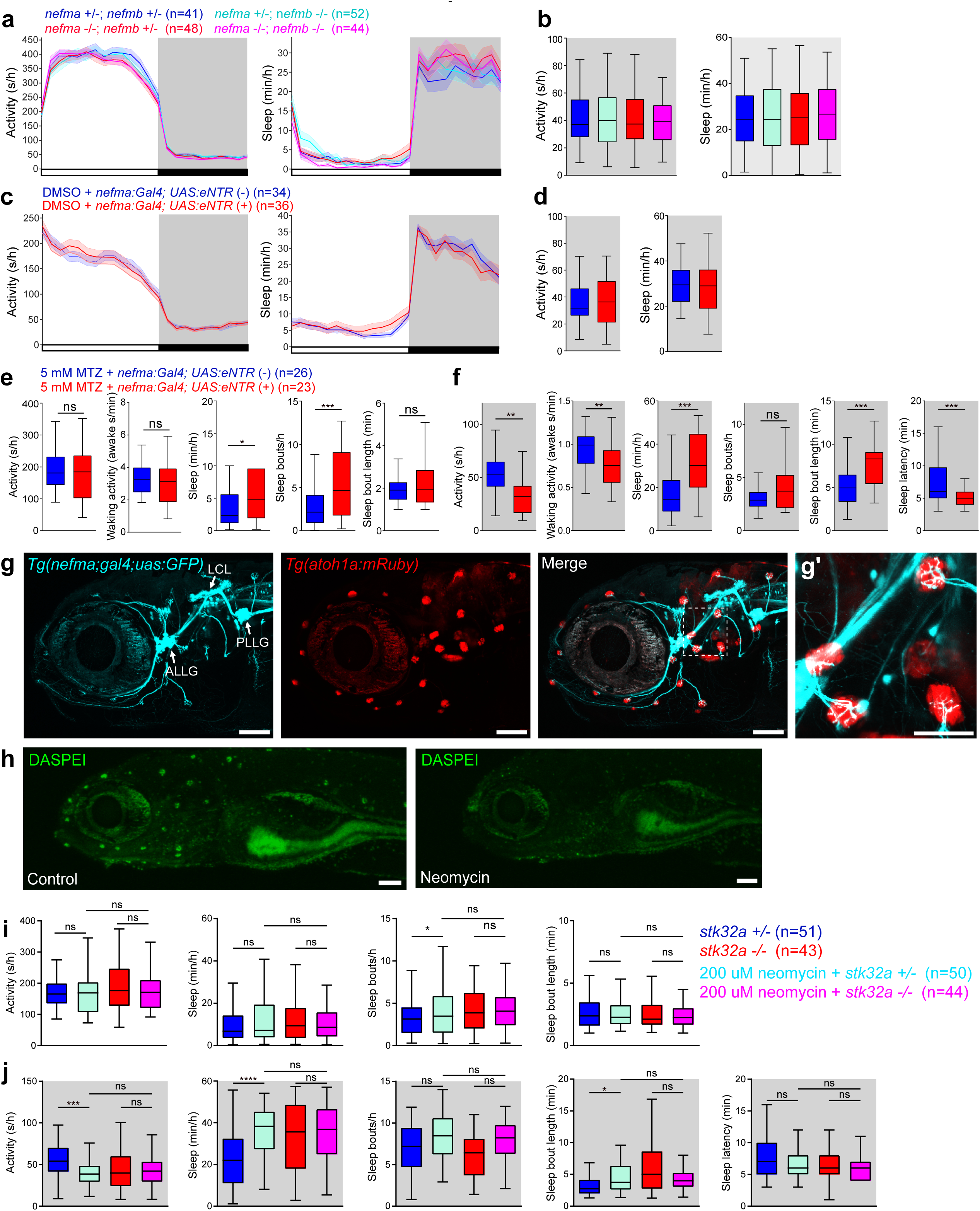
*nefma*-expressing neurons innervate lateral line neuromasts and ablation of neuromasts increases sleep. **(a,b)** Locomotor activity and sleep for *nefma* −/−; *nefmb*−/− zebrafish and their sibling controls at 6-dpf. **(c,d)** Locomotor activity and sleep for 6-dpf *Tg(nefma:Gal4); Tg(UAS:eNTR-mYFP)* and non-transgenic siblings treated with DMSO vehicle control. Genotypes indicate animals that were visually positive or negative for eNTR-mYFP fluorescence. Mean ± SEM **(a,c)** and box plots **(b,d)** are shown. **(e,f)** Box plots show locomotor activity, sleep, sleep bout number, and sleep bout length during the day **(e)** and night **(f)**, and sleep latency at night **(f)**, of 6-dpf *Tg(nefma:Gal4); Tg(UAS:eNTR-mYFP)* (red) and non-transgenic sibling control (blue) zebrafish treated with MTZ. Corresponding line graphs are shown in Figure 3E-3F. **(g)** Live imaging of a 6-dpf *Tg(nefma:Gal4); Tg(UAS:GFP); Tg(atoh1a:mRuby)* zebrafish shows that *nefma*-expressing neurons (cyan) in the ALLg and PLLg innervate neuromasts (red), and their afferent projections form the LCL in the brain (white arrows). Boxed region in the merged image is shown at higher magnification in **(g’)**. **(h)** Maximum intensity projections of live 6-dpf WT zebrafish treated with E3 vehicle control or 200 μM neomycin for 1 hour and stained with DASPEI shows loss of neuromast hair cells in neomycin-treated fish. **(i,j)** Box plots show locomotor activity, sleep, sleep bout number, and sleep bout length during the day **(i)** and night **(j)**, and sleep latency at night **(i)** of 6-dpf *stk32a +/−* and *stk32a −/−* zebrafish treated with 200 μM neomycin for 1 hour (cyan, magenta), or vehicle control (blue, red). Corresponding line graphs are shown in Figure 3G-3H. Black bars and gray shading indicate night. n = number of fish. ns, P>0.05; *P<0.05; **P<0.01; ***P<0.001; ****P<0.0001 by Mann-Whitney test **(d-f)** or Kruskal-Wallis test with Dunn’s test **(b, i, j)**. Scale bars: 100 μm **(g,h)**, 25 μm **(g’)**.

**Extended Data Figure 9.**
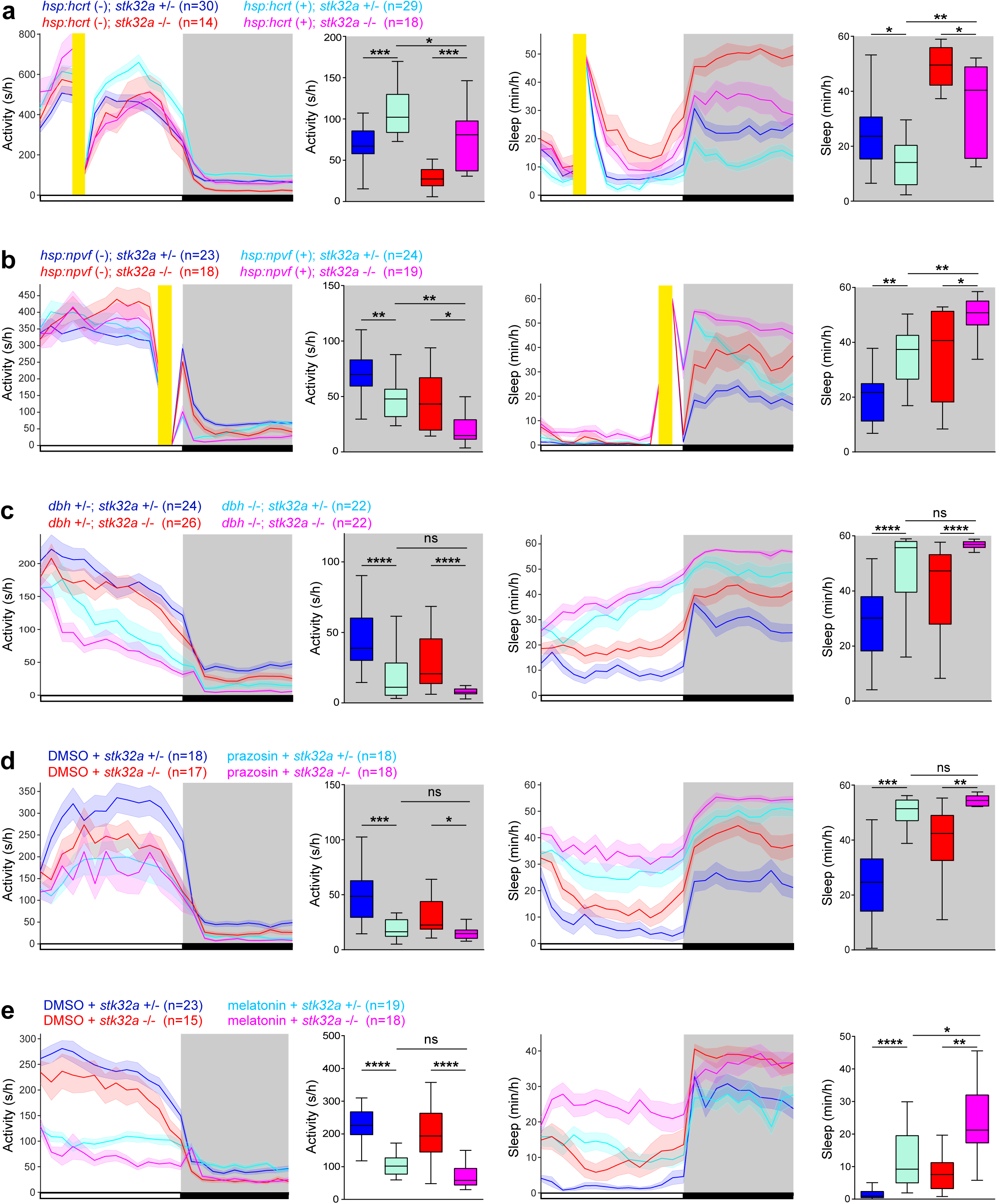
Epistasis experiments reveal genes and drugs that do not show interactions with *stk32a*. **(a)** Heat-shock induced overexpression of *hypocretin* (*hcrt*) in *Tg(hsp:hcrt)* fish results in more locomotor activity and less sleep at night in *stk32a* −/− fish compared to *stk32a* +/− fish. **(b)** Heat-shock induced overexpression of *neuropeptide vf* (*npvf*) in *Tg(hsp:npvf)* fish results in less locomotor activity and more sleep at night in *stk32a* −/− fish compared to *stk32a* +/− fish, and compared to *stk32a* −/− fish that do not overexpress *npvf*. **(c)** *dbh*−/−; *stk32a* −/− *fish* are less active and sleep more at night compared to *dbh* +/−; *stk32a* −/− fish. **(d)** Treating *stk32a* −/− fish with 2 μM prazosin (alpha1-adrenergic receptor inhibitor) results in less locomotor activity and more sleep at night compared to DMSO-treated *stk32a* −/− fish. **(e)** Treating *stk32a* −/− fish with 10 μM melatonin results in less locomotor activity and more sleep during the day compared to DMSO-treated *stk32a* −/− fish. Black bars and gray shading indicate night. Yellow bars indicate a 1-hour 37°C heat-shock. Mean ± SEM and box plots are shown. n = number of fish. ns, P>0.05; *P<0.05; **P<0.01; ***P<0.001; ****P<0.0001 by Kruskal-Wallis test with Dunn’s test.

**Extended Data Figure 10.**
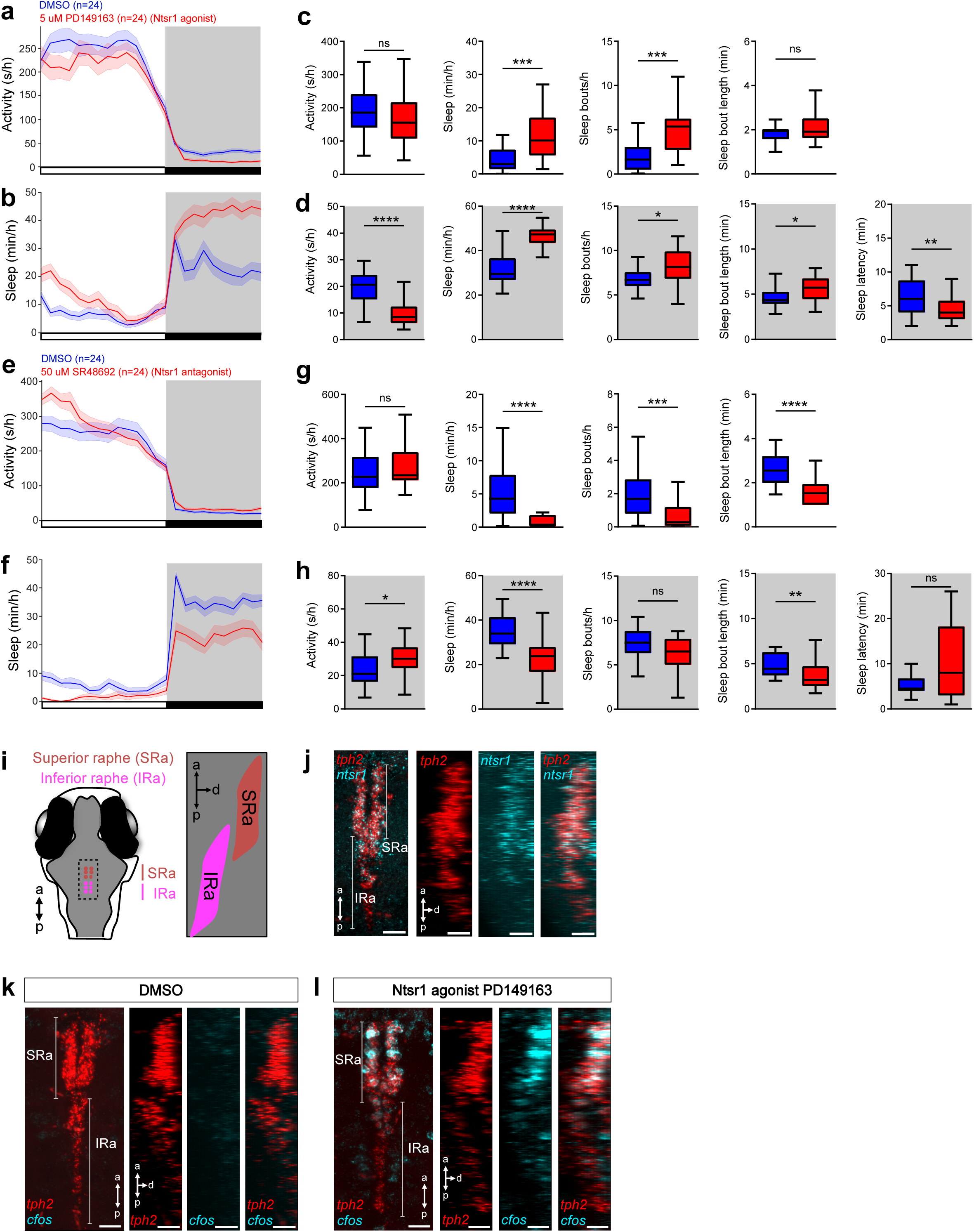
Effects of Ntsr1 agonist and antagonist on sleep and raphe neuron activity. **(a-d)** Treatment of 5-dpf WT zebrafish with 5 μM of Ntsr1 agonist PD149163 (red) resulted in less locomotor activity and more sleep at night compared to treatment with DMSO vehicle control (blue). **(e-h)** Treatment of 5-dpf WT zebrafish with 50 μM of Ntsr1 antagonist SR48692 (red) resulted in more activity and less sleep at night compared to treatment with DMSO vehicle control (blue). Black bars and gray shading indicate night. n = number of fish. ns, P>0.05; *P<0.05; **P<0.01; ***P<0.001; ****P<0.0001 by Mann-Whitney test. **(i)** Larval zebrafish brain schematic indicating locations of the superior raphe (SRa) and inferior raphe (IRa) (modified from ^56^). Dorsal (left) and side (right) views are shown. **(j)** HCR reveals that *ntsr1* is expressed in the SRa and the rostral inferior raphe IRa, which are marked by *tph2* expression. **(k,l)** Treatment with 5 μM of Ntsr1 agonist PD149163, but not with DMSO vehicle control, results in *c-fos* expression in the SRa and rostral IRa, similar to the expression *of ntsr1*. Side views in **(j-l)** were computationally generated by rotating z-stacks by 90° along the y-axis. Scale bars: 20 μm. a, anterior; p, posterior; d, dorsal.

**Extended Data Figure 11.**
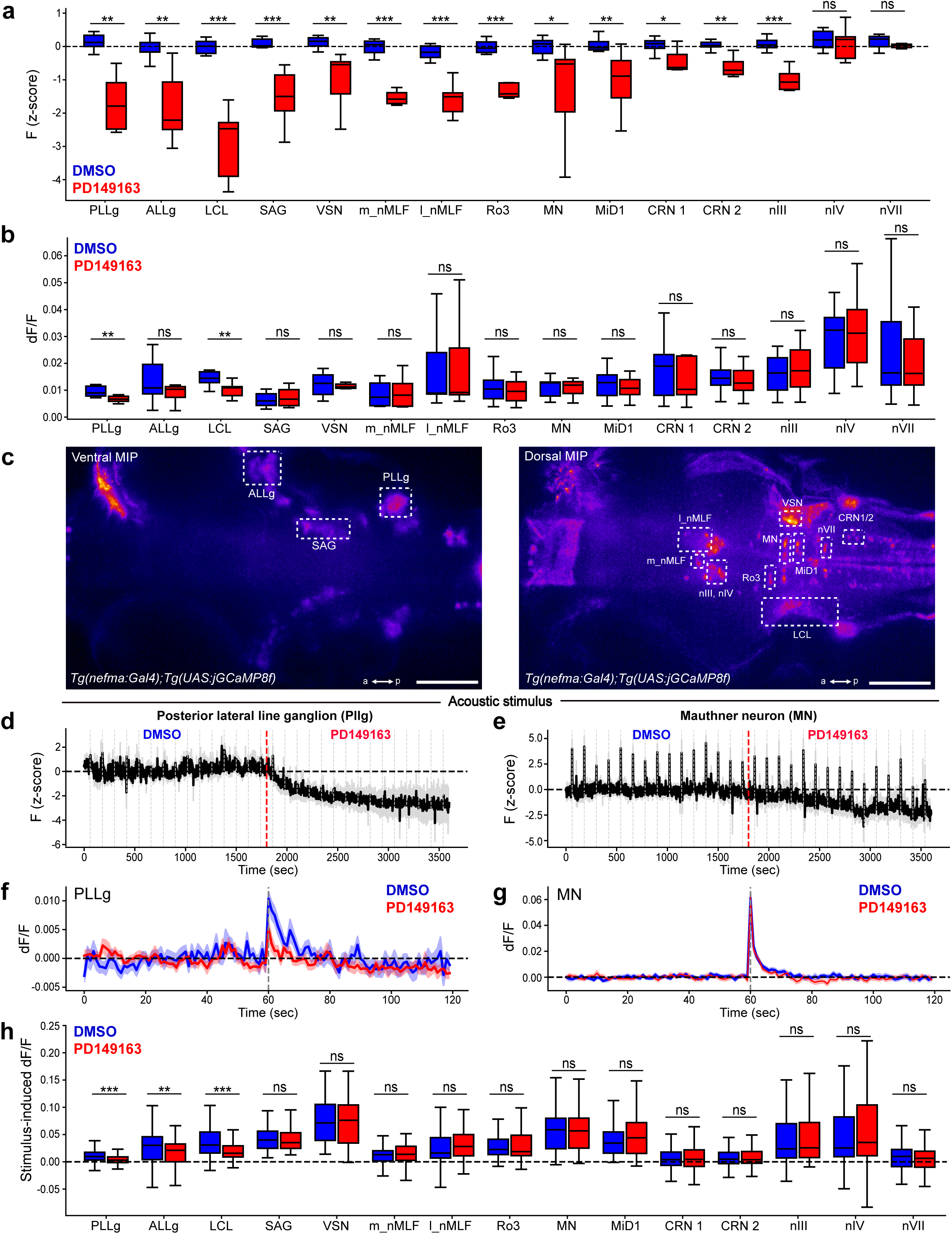
Ntsr1 signaling inhibits *nefma*-expressing sensory and motor populations. Box plots quantify raw fluorescence **(a)** and dF/F **(b)** recorded from *Tg(nefma:Gal4); Tg(UAS:jGCaMP8f)* fish in the regions indicated in **(c)** during 30 minute treatment with DMSO compared to 30 minute treatment with 5 μM of Ntsr1 agonist PD149163. **(c)** Maximum intensity projections (MIP) are shown for a 66 μm ventral stack (left) and a 66 μm dorsal stack (right) of the brain. a, anterior; p, posterior. **(d-g)** Acoustic stimulus-evoked jGCaMP8f fluorescence during treatment with DMSO and PD149163. Mean ± SEM raw fluorescence **(d,e)** and mean ± SEM dF/F for all acoustic stimulus trials with the stimulus applied at t=60 sec **(f,g)** for the posterior lateral line ganglion (PLLg) and Mauthner neurons (MN). Red dashed lines indicate start of PD149163 treatment. Gray dashed lines indicate acoustic stimulus presentations. Black dashed lines indicate y=0. **(h)** Box plots compare jGCaMP8f dF/F responses to the acoustic stimulus at t=60 sec, normalized to baseline (t=0-59 sec) of each trial, for DMSO and PD149163-treated fish. n = 7 fish. ns, P>0.05; *P<0.05; **P<0.01; ***P<0.001 by Mann-Whitney test.

